# Sla2 is a core interaction hub for Clathrin Light Chain and the Pan1/End3/Sla1 Complex

**DOI:** 10.1101/2024.11.14.623549

**Authors:** George Draper-Barr, Lucas A. Defelipe, David Ruiz-Carrillo, Emil Gustavsson, Meytal Landau, Maria García-Alai

## Abstract

The interaction network of Sla2, a vital adaptor protein in the endocytic mid-coat, undergoes constant rearrangement incorporating or replacing interacting proteins over time. Sla2 serves as a scaffold linking the membrane to the actin cytoskeleton, with this role modulated by Clathrin Light Chain (CLC), which inhibits Sla2’s function under certain conditions. We show that Sla2 has two independent binding sites for CLC: one previously described in homologs of Fungi (Sla2) and Metazoa (Hip1R), and a second found only in Fungi. We present the structural model of the Sla2 actin-binding domains in the context of regulatory structural domains by electron cryo-microscopy. We provide an interaction map of Sla2 and the regulatory proteins Sla1 and Pan1, predicted by AI modelling and confirmed by molecular biophysics techniques. Pan1 competes with CLC for the conserved binding site on Sla2. These results enhance the mapping of crucial interactions at endocytic checkpoints and highlight the divergence between Metazoa and Fungi in this vital process.

**Teaser:** Sla2 forms complexes with three regulatory proteins in the endocytic pit, two of which compete for the same site

## Introduction

Endocytosis is the essential process of macromolecule internalisation from the extracellular space, and recycling of membrane receptors through transport to internal membrane systems by packaging cargo into membrane-bound vesicles. This process has evolved during the course of eukaryogenesis, establishing isolated transport events and contributing to the diversity of cellular activities within internal compartments (*1–3*). This system relies on a host of protein families that have diverged and duplicated over the course of time (*1*, *2*, *4*). Research on clathrin-mediated endocytosis (CME) in budding yeast has provided a detailed understanding of its timeline and components, involving over 60 proteins through genetic, microscopy, and structural studies (*5–8*). The stages of endocytosis require the precise timing of cargo recognition, membrane curvature-inducing proteins, scission proteins, and GTPases (*9–11*). Based on their time of arrival at the endocytic pit, proteins have been categorised into different modules: early arriving proteins (e.g. Ede1), mid-coat proteins (Sla2 and Ent1/2), coat proteins (e.g. Clathrin), WASP/Myo proteins, actin module proteins, and scission proteins (e.g. Dynamin) (*7*). The work described in this study focuses on Sla2 (homologous to *Hs*Hip1R) and its regulatory role in the mid-coat of the endocytic pit.

The N-terminal region of Sla2 contains an ANTH (AP180 N-Terminal Homology) domain, which anchors Sla2 to the plasma membrane by forming complexes with PI(4,5)P2 and the ENTH (Epsin N-Terminal Homology) domains of Ent1/2 (Figure 1) (*12*). The AENTH complex formed between Ent1/2 and Sla2 is crucial for recruiting Sla2 to the endocytic pit, as mutations in this domain significantly reduce cargo internalisation *in vivo* (*13*). This pair of proteins also interact directly to the Clathrin coat of the endocytic pit. Ent1 binds to the 7-blade β-propeller domain of Clathrin Heavy Chain via a Short Linear Motif (SLiM) at the C-terminus of its Intrinsically Disordered Region (IDR) (*14*), and Sla2 through its coiled-coil to the N-terminal Intrinsically Disordered Region (IDR) of Clathrin Light Chain (*15–19*). The Clathrin heterodimer, comprising the heavy (CHC) and light (CLC) chains, forms a ‘triskelion’ structure that assembles into a lattice covering the surface of the endocytic pit (*20*). The Clathrin Light Chain (CLC) has three segments: an N-terminal IDR that binds CHC and Sla2, a central helix that interacts with CHC’s distal leg, and a C-terminal region thought to bind CHC’s trimerization domain (*21*, *22*).

**Figure 1:**
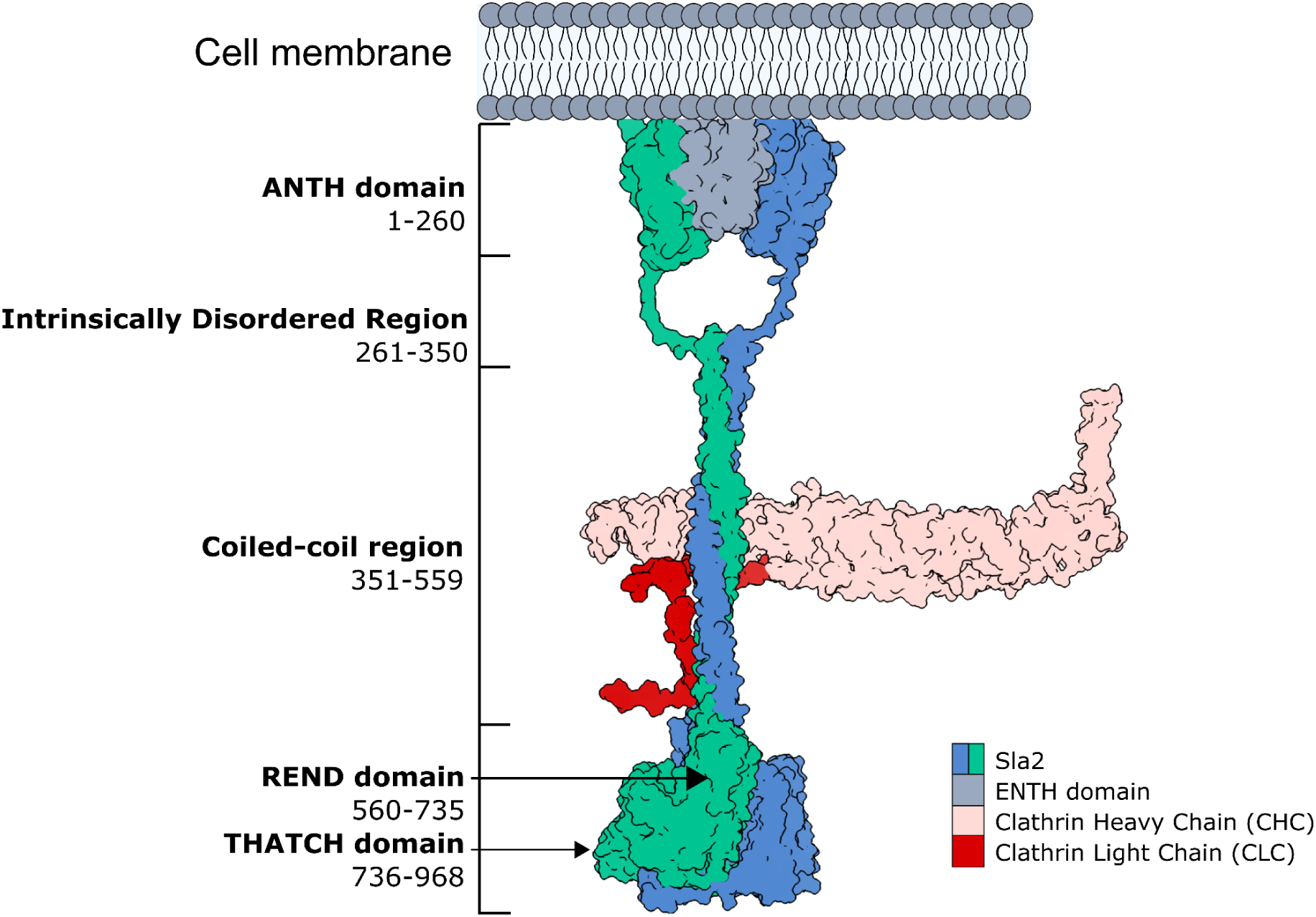
Sla2 domain architecture in complex with the Clathrin heterodimeric coat. The primary complexes of interest involve Sla2, Ent1, and the Clathrin triskelia. The ANTH domain of Sla2 covers residues 1-260, IDP region from 261-350, coiled-coil from 351-559, REND domain from 560-735, and THATCH domain from 736-968.

Sla2 contains an Intrinsically Disordered Region (IDR; residues 261-350) and a coiled-coil segment (residues 351-559) between the ANTH domain (residues 1-260) and the C-terminal region (residues 560-968). One of the other important roles of Sla2 in the endocytic pit is the regulation of the Actin polymerisation complex Pan1/End3/Sla1. This complex modulates Arp2/3 and Las17 activity that in turn regulates endocytosis in yeast, shown by depletion experiments from the work of Sun et al. (*23*, *24*). The removal of the Sla2 interactions lead to erroneous Actin regulation and subsequent decrease in processive endocytic events. Sla2 forms separate complexes with Sla1 and the Pan1 coiled-coil, these interactions cause the release of Las17 from Sla1 and inhibits positive regulation of Actin polymerisation by Pan1 (*25*, *26*). The current domain architecture of Sla1 includes three SH3 domains (two of which bind Las17), the Sla1 Homology Domain (SHD1), and a SAM domain (SHD2). SH3 domains primarily bind proline rich motifs (*27–29*). The Sla1 Homology Domain (SHD1) is responsible for binding NPFxD motifs in endocytic cargo (*30*). The SAM domain (SHD2) negatively regulates Sla1 binding the Clathrin Heavy Chain via its variant Clathrin Binding Motif in the C-terminus (*31*). The exact location and quantitative measurement of each interaction with Sla2 is unknown for both proteins.

The Actin-binding domain from Sla2/Hip1R is named the Talin-Hip1/R/Sla2p Actin-tethering C-terminal homology (THATCH) domain, with critical residues for Actin binding on the termini of helices 3 and 4 of the THATCH core, determined through mutagenesis and Actin co-sedimentation assays (*32–34*). Cell biology studies have localised the Actin-binding domains of *Hs*Hip1R in proximity to Actin filaments at the endocytic site (*35*). This interaction is in combination with Ent1 that contains a C-terminal Actin Binding Domain (ABD) (*36*). The ABD of Ent1 and C-terminus of Sla2 are redundancies for each other in the total connection network of the membrane to actin filaments. Mutational studies of the Actin binding domains of Ent1 and Sla2 show a significant decrease in internalisation only from a double mutant of both domains. This led to the proposal that Sla2 and Ent1/2 form a large island of protein motifs for recruitment of Clathrin and Actin. This would organise an Actin network over the plasma membrane surface and the relaxation of the Clathrin triskelia. The THATCH domain is formed of two parts: the THATCH core that binds Actin, and the LATCH helix that is a key dimerisation motif of the C-terminal region (*33*). Removing either portion stops the binding of Actin. The REND domain, a force sensing domain described in End4 (homologous to Sla2) has been compared to the R12 domain of Talin (*37–39*). This domain is N-terminal of the THATCH domain, and unfolds above a force threshold that is exerted by Actin during endocytosis. This unfolding process eliminates the Actin binding of Sla2 as well, suggesting that the REND domain acts as a stabilising and a regulatory factor of the C-terminal region structure during endocytosis. The arrangement of the REND domain, the THATCH domain core and the LATCH helices in the complete structure of this region is unknown. Answering this would determine the availability of Clathrin regulated Actin binding surface of the THATCH domain, as well as the relationship of the conformational changes of the THATCH and REND domains to expose this surface.

Hip1R and its relative, Hip1, have been shown to interact with Clathrin Light Chain (*16*, *18*, *40–45*). This interaction occurs in the coiled-coil region of the adaptor, with a specific region proposed from the crystal structure of the Hip1 coiled-coil, forming a platform for CLC binding (*15*, *16*, *46*). In yeast and mammals, the interaction between CLC and Sla2/Hip1R inhibits Actin binding by the THATCH domain, as well as regulating the Clathrin lattice stiffness (*16*, *17*, *32*, *33*, *47*). The exact location, magnitude, and function of the CLC interaction with Sla2 for the processivity of endocytosis was not previously described. This is particularly important as the endocytic system between Fungi and Metazoa differs in the critical role Actin takes in endocytosis of Fungi as opposed to Metazoa.

In this study, we characterise the biophysical properties of the Sla2-CLC complex and identify a novel binding site unique to Fungi, located near a site conserved across both Metazoa and Fungi. Our work also shows that these binding sites are independent of each other *in vitro*. We determined the structure of the C-terminal region of Sla2 by electron cryo-microscopy (cryo-EM). This allowed us to put prior information on the Actin binding role of this region into three-dimensional context with the regulatory REND domain and LATCH helix. We have also characterised the endocytic dynamics of CLC binding site mutants *in vivo* in contrast to the WildType protein. In addition, we have shown that the coiled-coil of Sla2 and Pan1 complex together with a similar affinity as the CLC interaction and that the IDR of Sla2 is responsible for interactions with Sla1. This work has furthered the understanding of the highly regulated endocytic process in *S. cerevisiae* and show that there is a divergence in the sequence of Sla2 as compared to Hip1R that can be directly related to the new interactions we shed light on.

## Results

### Sla2 forms complexes, via the coiled-coil, with CLC through two independent sites

Clathrin Light Chain interacts with Sla2/Hip1R across both mammalian and fungal systems (*16*, *17*). This interaction in the human system was shown to be mediated by the coiled-coil of Hip1R and the disordered N-terminus of CLC. In order to characterise this interaction we used a construct of residues 351-968 of Sla2 (Sla2ccRTH) and NusA conjugated to CLC (NusA-CLC) (Figure 2a). With these constructs we performed Mass Photometry (MP) experiments (*48*, *49*) that allowed us to identify individual masses of high affinity complexes with heterogeneous stoichiometry (*50*, *51*). We measured samples at 50 nM of each component, Sla2ccRTH and NusA-CLC, and in combination with each other. The results at these concentrations showed that the Sla2 construct is dimeric and the NusA-CLC is monomeric (Supplementary Figure 1). In the mixture of Sla2ccRTH and NusA-CLC we observed peaks for the individual components as well as two other peaks, which correspond to one dimer of Sla2 bound by one NusA-CLC and a large complex of undetermined stoichiometry (Figure 2b). The mixture of NusA and Sla2ccRTH showed no increase in mass so we can eliminate the effect of conjugating NusA to CLC on complex formation (Supplementary Figure 1).

**Figure 2:**
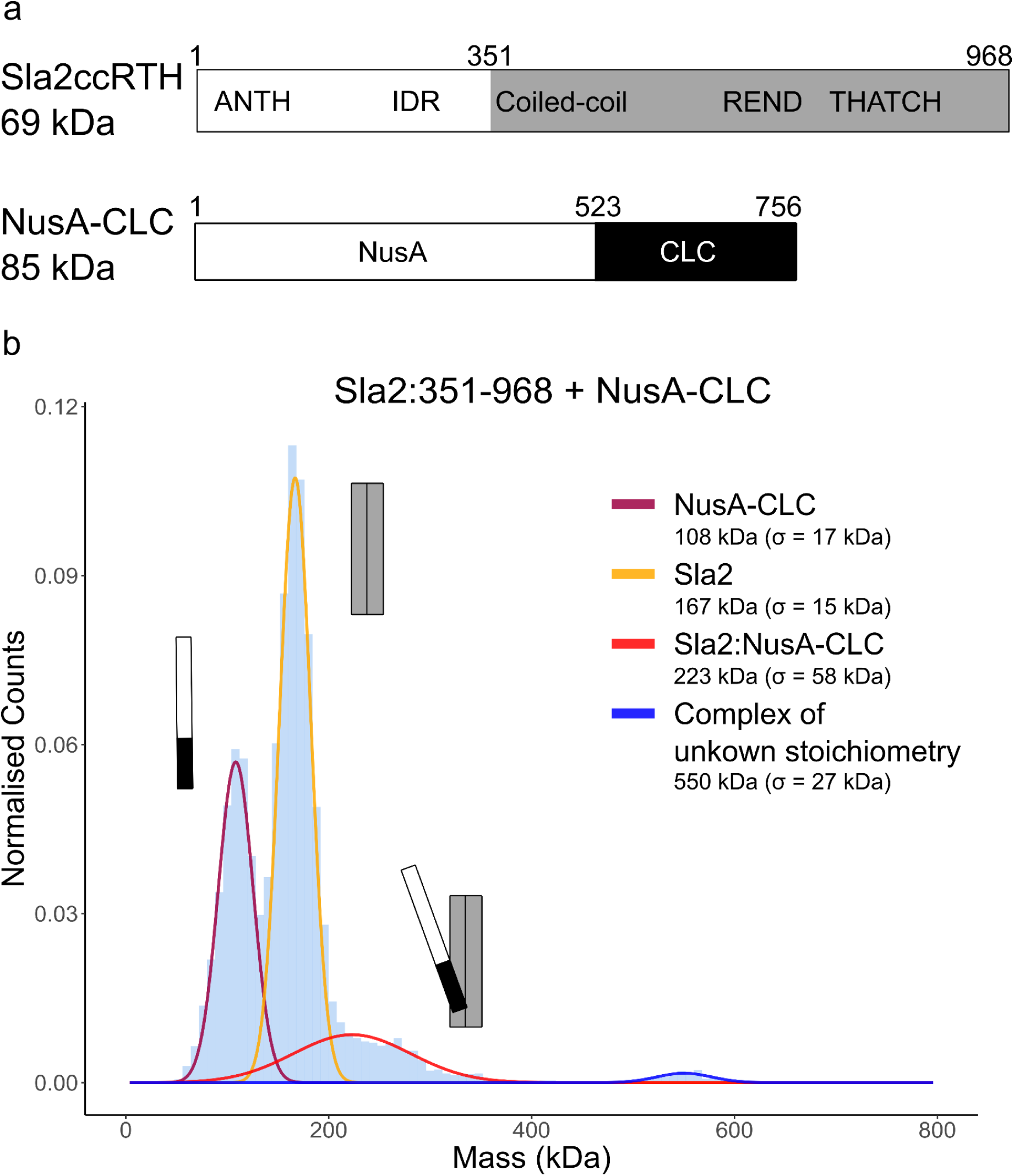
Mass Photometry captures the Sla2:CLC complex at sub-micromolar concentrations (**A**) Sla2ccRTH construct explained in context of the full length protein, NusA-CLC construct diagram showing the N-terminal moiety of a fusion protein, NusA, to the CLC full length polypeptide. (**B**) Mass Photometry measurements of Sla2:351-968 (Sla2ccRTH) and NusA-CLC. Two populations of particles measuring at masses of 108 kDa (σ = 17 kDa, 31 % counts, NusA-CLC), and 167 kDa (σ = 15 kDa, 52 % counts, Sla2ccRTH). Larger particles corresponding to a mass of 223 kDa (σ = 58 kDa, 15 % counts, Sla2ccRTH+NusA-CLC), and 550 kDa (σ = 27 kDa, 1 % counts) are measured. Gaussian fitting of Refeyn 2.0 exported events achieved through the eSPC program, PhotoMol (*51*).

In order to understand the binding of CLC and the Sla2 coiled-coil further, we determined the affinity of this complex using Microscale Thermophoresis (MST) (*52*). The titration of Sla2cc (Sla2 residues 296-767) against CLC showed two separate binding events with different Kds (Figure 3). The value for the lower affinity site (Site 2) was determined as 3.0 μM (CI95: 2.07-4.29 μM) and the higher affinity site (Site 1) at ∼100 nM across the three Sla2cc constructs tested. The affinity of Site 2 is in agreement with the value previously reported for this complex in the human homolog (*16*, *45*). To determine which portions of the Clathrin Light Chain are responsible for different sites, we titrated shortened constructs of CLC against Sla2cc. From these experiments we have identified Site 1 to be located in residues 70-140 of the CLC N-terminus, and Site 2 is located in residues 1-70 which corresponds to the Intrinsically Disordered Region also proposed in the human homolog (Supplementary Figure 2).

**Figure 3:**
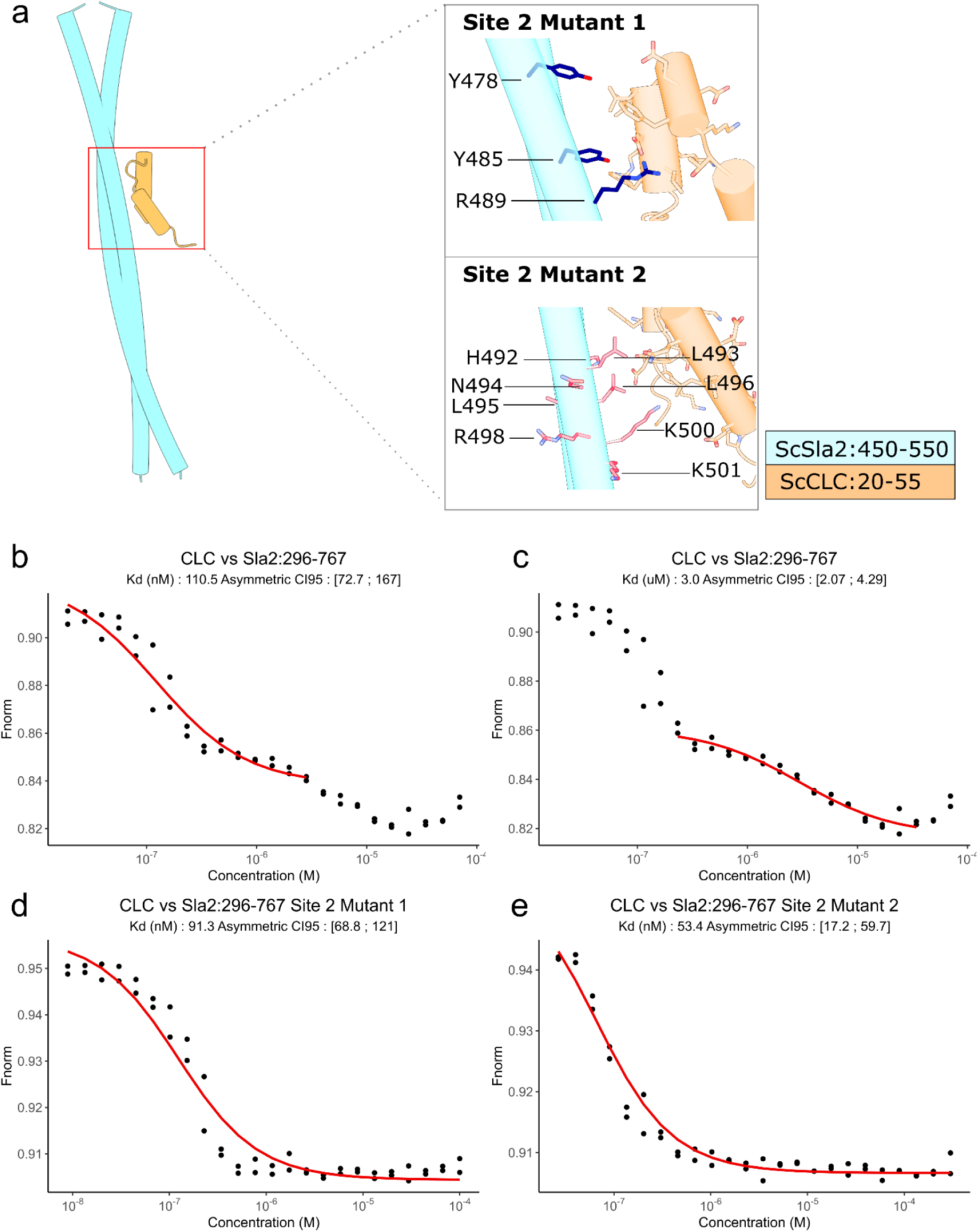
Sla2 has two independent binding sites for CLC (**A**) AF3 model of the Sla2:CLC complex for residues 450-550 of dimeric Sla2 and 20-55 of CLC. Highlighted in the insets are the residues of the coiled-coil for one chain that are mutated in each construct used to map Site 2, dark blue for Site 2 Mutant 1 and red for Site 2 Mutant 2. (**B**-**C**) Fitted MST curves for the WildType *Sla2* against labelled CLC. There are two separate binding events. (**B**) The high affinity event was measured at ∼100 nM across the three constructs. (**C**) The low affinity event was measured at 3.0 μM (CI95: [2.07, 4.29]) solely in the Wild Type coiled-coil construct. (**D**-**E**) Two separate mutants of Sla2 were determined to abrogate binding for the lower affinity binding site of CLC. (**D**) Sla2cc Site 2 Mutant 1 (Y478A, Y485A, R489A), and (**E**) Sla2cc Site 2 Mutant 2 (H492A, L493A, N494R, L495A, L496A, R498G, K500D, K501E, L502A). Both mutants only have one transition, which corresponds to Site 1 by the value of the K_d_.

To determine which residues were required for the different binding events to Sla2cc and if the sites are independent of each other, mutations were done on each site. For Site 2, two mutants were generated from conserved residues in this region between Metazoa and Fungi (Figure 3a). For Site 2 Mutant 1 we used the following mutations: Y478A, Y485A, R489A. In the case of Site 2 Mutant 2 we mutated the following residues: H492A, L493A, N494R, L495A, L496A, R498G, K500D, K501E, L502A. Both mutants eliminated Site 2 independently of Site 1 (Figure 3d-e). These mutants highlight the sensitivity and broad surface area required for this interaction site, which contains numerous hydrophobic and charged residues. Cross-Linking-MassSpectrometry experiments were also a key tool in mapping Site 2 and validated the AlphaFold3 model of the Sla2:CLC complex that could also predict the Site 2 interaction (*53*). These BS3 cross-linking experiments between Sla2cc and CLC:1-80 gave several intermolecular cross-links, residues 20, 21, and 29 of CLC cross-linked to Sla2 residue 477, and a cross-link between residues 55 of CLC and 505 of Sla2 (Supplementary Figure 3).

Alignment of the cross-linked regions from both proteins shows high conservation of this region across Fungi and Metazoa, indicating its likely critical functional role (Supplementary Figure 3). Coiled-coil residues are exposed in the cleft in the AlphaFold3 model of Sla2 for potential interactions to the N-terminus of the CLC. Coordinating residues Y478, Y485, and R489 of Sla2 are residues 28 to 44 of CLC, particularly charged residues E31, E38, and D44. CLC F39 is in proximity to Sla2 Y485. In the proposed Hip1 motif region, H492 is contacting F27 of CLC and N494 with Q43. It is of note that L493 is in proximity with L28 and L47 of CLC in our AF3 models (Figure 3a). The Sla2 coiled-coil relies on aliphatic residues, with modelling suggesting that these residues guide the Clathrin Light Chain (CLC) into the coiled-coil groove for stable binding.

The molecular identity of Site 1 needed to be understood as from the results until this point we could only determine that the CLC residues 70-140 were responsible for this interaction and that Site 1 is within the coiled-coil of Sla2 but not in the region of residues 478-505. We analysed conservation of the coiled-coil across both Fungi and Metazoa. We observed that the region C-terminal of Site 2 in the coiled-coil contains significant conservation that is only present in Fungi (Figure 4a). There is no structural information available for the interaction of Sla2 and CLC in this region. The AlphaFold3 model has no confidence for an interaction between them either. A deletion mutant of residues 515-546, Sla2cc ΔSite1 (Δ515-546), was devised to remove Site 1 from Sla2. We have used Circular Dichroism to determine the secondary structure and thermal stability of this construct and the other constructs used for MST determination of K_d_ measurements (Supplementary Figure 4). The construct was folded with a very similar helical content compared to the WildType sequence. Using this construct then to measure MST against labelled CLC gave only one transition with a K_d_ value in the micromolar range, which corresponds to Site 2 (Figure 4b). From this result we can map Site 1 to residues 515-546 of Sla2 (Figure 4c) and conclude that this site is a specific interaction not seen in Metazoa through sequence conservation.

**Figure 4:**
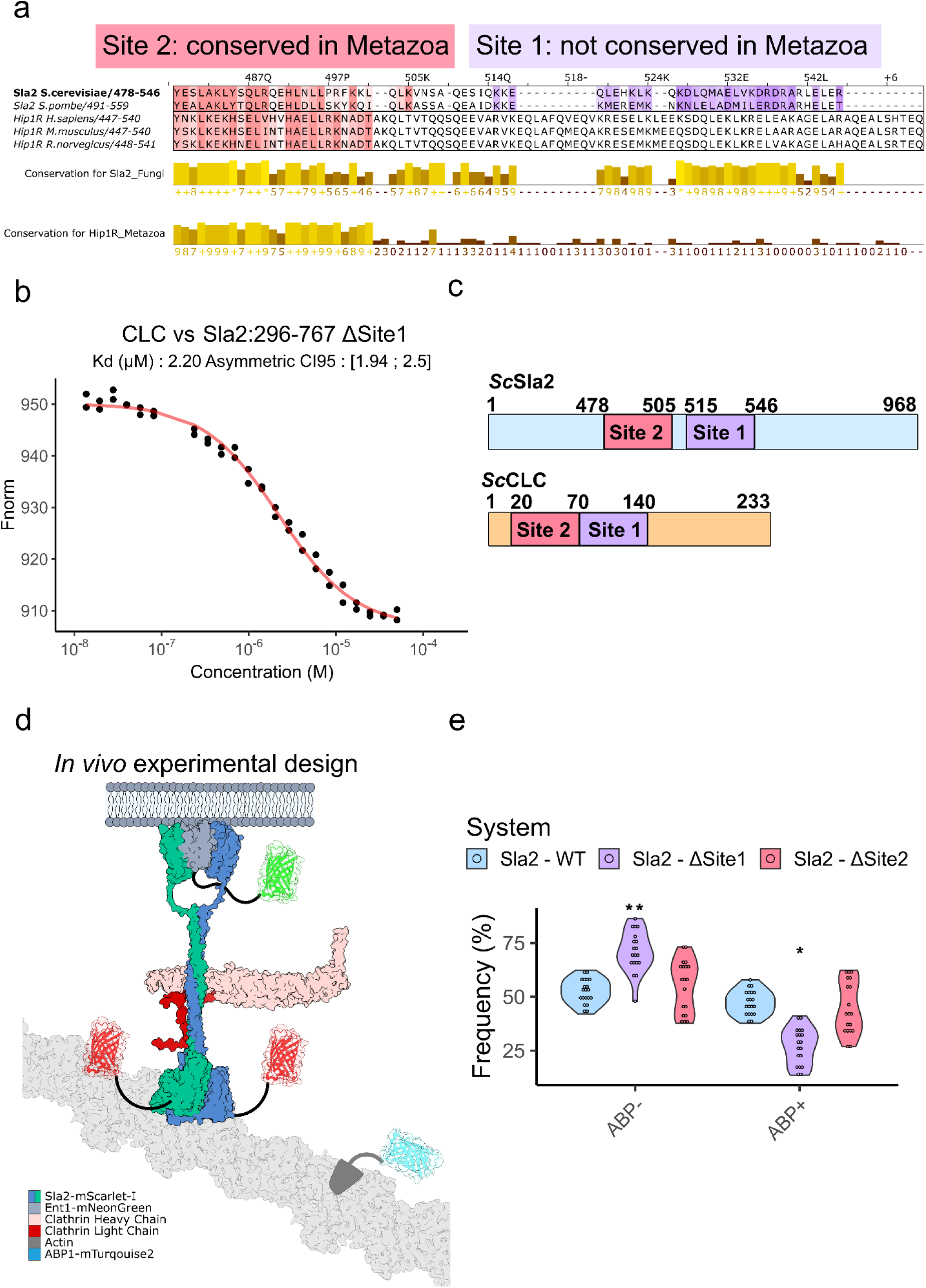
Fungi have a second interaction site between Sla2 and CLC (**A**) Sequence alignments of the coiled-coil region of Sla2 and Hip1R in both Fungi and Metazoa, with representative model species shown in the figure. Site 2 (residues 478 to 505) is coloured in red with above 30 % conservation, the proposed Site 1 region between residues 515 and 546 of *Sla2* and the aligned regions of the other sequences is also highlighted in purple above 30 % conservation. We can observe that there is significant conservation in the Site 1 region of Fungi not seen in Metazoa. (**B**) MicroScale Thermophoresis of Sla2cc ΔSite1 (Δ515-546) titrated against CLC (REDNHS labelled). (**C**) A diagram of Sla2 and CLC and the locations of the binding sites between these two proteins. The high affinity interaction (Site 1) between Sla2 and CLC is found between residues 515 and 546 of the coiled-coil and Site 2 is located further towards the N-terminus of Sla2 at residues 478 to 505. (**D**) Endocytic dynamics was measurable using an endogenously tagged Ent-1-mNeonGreen, Abp1-mTurqouise2 *S. cerevisiae* strain with ectopically expressed Sla2-mScarlet-I under an endogenous promoter. (**E**) Abp1 positive and negative events determined by cmeAnalysis (*56*) for Sla2 WT, ΔSite1, and ΔSite2 (ΔYYR). ΔSite1 is significantly reduced in the percentage of Abp1+ events as compared to WT and also ΔSite2. Asterisks indicate statistically significant differences (** - p = 5.8e^-5^, * - p = 0.00026). Two-sided Mann-Whitney U-test corrected with Benjamini and Hochberg for multiple comparisons was used to calculate p values.

In order to discern if the two sites are important to the normal function of endocytosis, *in vivo* endocytic dynamics were measured within *S. cerevisiae*. For tracking the progress of endocytosis, Abp1 and Ent1 were endogenously tagged with mTurqouise2 and mNeonGreen, respectively, as previously done by Defelipe et al. (*54*) (Figure 4d). Sla2 is expressed ectopically in a vector with the native promoter with knocked-out endogenous Sla2, using WildType, Site 2 Mutant 1 (ΔSite2), and ΔSite1 Sla2 sequences (Figure 4e). Abp1 is involved in the recruitment of Actin (necessary for membrane invagination and vesicle formation) and therefore is associated with a productive endocytic event while its absence towards the end of Ent1 lifetime would represent an abortive site (*55*). We classified endocytic events into two main categories (using the program cmeAnalysis(*56*)): Ent1 and Abp1 positive (Abp1+) or Ent1 positive and Abp1 negative (Abp1-) (Supplementary Figure 5). From our analysis of the endocytic events measured using TIRF microscopy, the ΔSite1 mutant of Sla2 causes a significant decrease in Abp1+ events, ΔSite2 has no such distinction from the WildType (Figure 4e).

### Structure determination of the Sla2 C-terminal region

The experimental structure of the REND domain and the arrangement of the THATCH domain to this domain is unknown. The arrangement of the THATCH domain is important when it comes to understanding the Actin binding mechanism of Sla2 as well. To obtain answers to these questions surrounding a key interaction in fungal endocytosis, we used cryo-EM to structurally characterise the C-terminal regions of Sla2. The construct Sla2ccRTH was used again as this contains the full coiled-coil and C-terminal region of Sla2. Sla2ccRTH was imaged and subsequent processing resulted in a density map at a resolution of 3.6 Å. The resolution is reported from the FSC_(0.143)_ resolution determination from Refmac (*57*). This resolution allows a model for the map to be fitted from a base model from the AlphaFold3 predictions of the Sla2 domains (Figure 5). The mean RMSD of the AF3 model to the experimental model is 2 Å. The complete C-terminal region (REND and THATCH domains) had not previously been determined experimentally, although the individual THATCH core of *Hs*Hip1R has been crystallised (Figure 6 and Supplementary Figure 6a). The high similarity to the AlphaFold model made the process significantly faster as it allowed us to use a confidently folded model for refinement without a high resolution map. This convergence of the experimentally determined and computationally determined models for this sequence highlights the ability of AlphaFold to predict previously unseen combinations of domains that are then experimentally confirmed. Our results as well show a highly similar structure to the *Hs*Hip1R THATCH core. Due to the high sequence and structural conservation with Hip1R and mutational studies on the THATCH core (*32*) we can highlight in our structure where the Actin binding region of Sla2 is located. The structure of the complete C-terminus of Sla2 shows that the proposed Actin binding surface is not fully available to the solvent and instead has several contacts to the REND domains (Supplementary Figure 6b).

**Figure 5:**
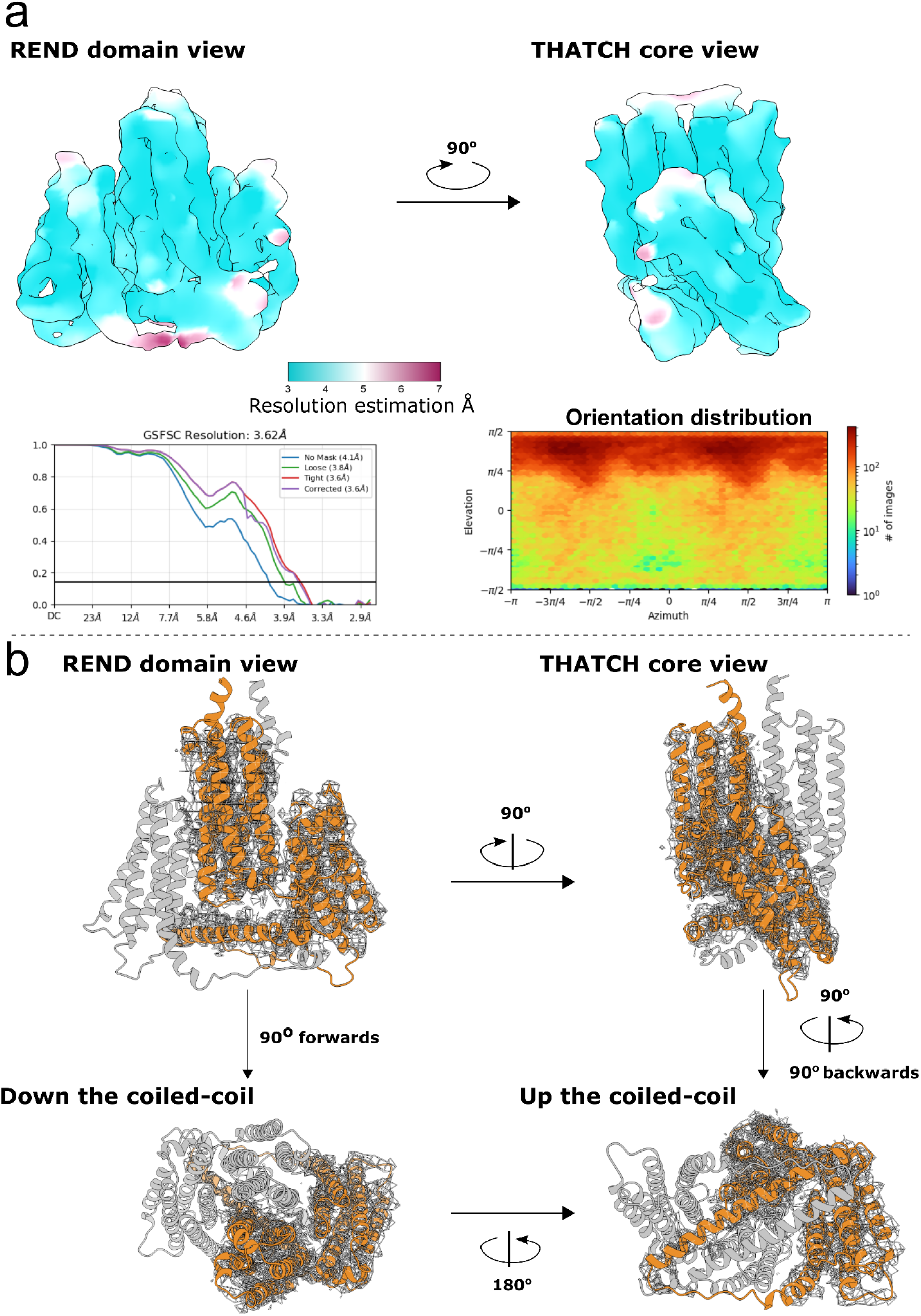
Sla2 C-terminal domains resolved below 4Å (**A**) Density maps for the Sla2ccRTH construct density map. The maps are colour coded according to the local resolution estimation by cryoSPARC v4. There are two graphs present as well for the density map, on the bottom left, the GSFSC Resolution graph determined post auto-tightening of the map and, bottom right, the orientation distribution map of the particles. Estimation of the resolution through Refmac is 3.62 Å. (**B**) Sla2 residues 560 to 968 are modelled into the locally filtered density map shown for one chain, obtained from micrographs collected on Sla2ccRTH. This construct contains all of the coiled-coil region as well as the C-terminal domains, however we modelled the REND domain through to the end of the LATCH helix.

**Figure 6:**
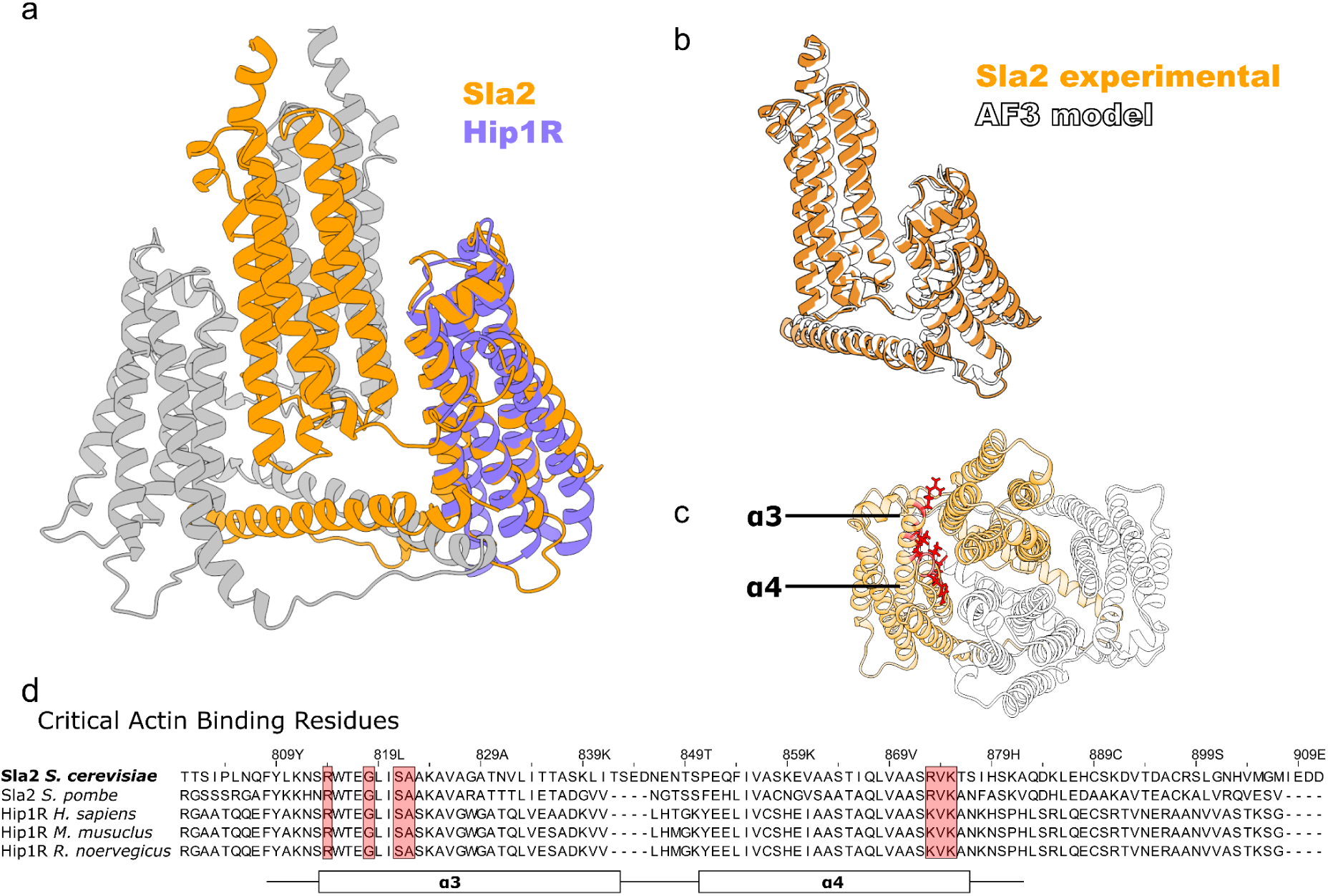
Actin binding residues conserved between Sla2 and Hip1R have altered intrachain interactions between conformations (**A**) Overlay of the ScSla2 560-968 model with the *Hs*Hip1R THATCH core crystal structure (PDB:1r0d) matched to Chain A of our structure. The overall RMSD of 1r0d to the residues 735-909 of Sla2 is 6.3 Å, however the overall fold is the same and the RMSD of the Actin binding residues is 0.8 Å. (**B**) Overlay of one chain of both the Sla2 experimental model and the AF3 computational model for residues 560-968 of Sla2. The RMSD of the chains is 2 Å. (**C**) The view of our model from the perspective of looking down the coiled-coil. Highlighted are residues that are conserved and considered critical for binding Actin fibrils in red. These residues are in contact with the REND domain interface. (**D**) Alignments of helices 3 and 4 of the THATCH domain for the Fungi Sla2/End4 and Metazoa Hip1R sequences with representative species shown. *Hs*Hip1R THATCH domain studies showed residues in the N- and C-termini of ɑ3 and ɑ4 respectively are critical to Actin binding (*32*). These residues are highlighted in red.

The REND domain in our structure is folded into a dimeric 5 helical bundle and the LATCH helices form an antiparallel dimer with the respective C-termini position towards the THATCH core domain of its own chain (Supplementary Figure 7a). The N-terminus is positioned away from the THATCH core of its respective Sla2 chain next to helix 3 of the other chains THATCH core and then passes under the REND domain before the C-terminus reaches back to its own THATCH core helix 3. The proposed dimerisation surface from Talin-1 (*33*) that was applied to Sla2 and Hip1R is not consistent with our model (Supplementary Figure 7b). The LATCH helix structure starts at 925 through to 968 and isn’t aligned between residues Q837 to Y962. The THATCH core interface with the REND domain face contains helices 3-5 of the THATCH domain. The buried solvent accessible area between the LATCH helices and the REND domains of the Sla2 C-terminal structures is 65 Å^2^. The LATCH helix is shifted down from the core domain and has a space between the REND domain and LATCH helix (Supplementary Figure 7c). The buried surface area between the THATCH domain of a single chain with the REND domains of both chains is 750 Å^2^.

### Sla2 forms two separate interactions in distant locations with the Pan1/End3/Sla1 regulatory complex

As introduced earlier, Sla2 forms a complex with Sla1 as part of the disruption of the Pan1/End3/Sla1 complex to release Las17 (*25*, *26*, *58*, *59*). Using AlphaFold3 (AF3) (*53*), we modelled Sla1 residues 120 to 510, which is the region between SH3_2 and SHD1 in the Sla1 annotated structure (Figure 7a). In the AF3 prediction, between the 2nd and 3rd SH3 domains, there is a well-predicted, unannotated folded domain, whose highest-ranking structurally homologous domains from FoldSeek (*60*) belong to the Pleckstrin Homology (PH) domain family (*61*). These domains bind PIPs, one of the highest-ranked hits was the 2nd OPY1 PH domain (*62*), which specifically binds PI(4,5)P_2_ (Supplementary Figure 8). In addition, searching FoldSeek using the AF3 model of the 2nd OPY1 PH domain gave the Sla1 PH domain as a confident hit. The only other high confidence prediction domain in the Sla1:120-510 AF3 model is the previously annotated 3rd Sla1 SH3 domain (SH3_3) (Figure 7a).

**Figure 7:**
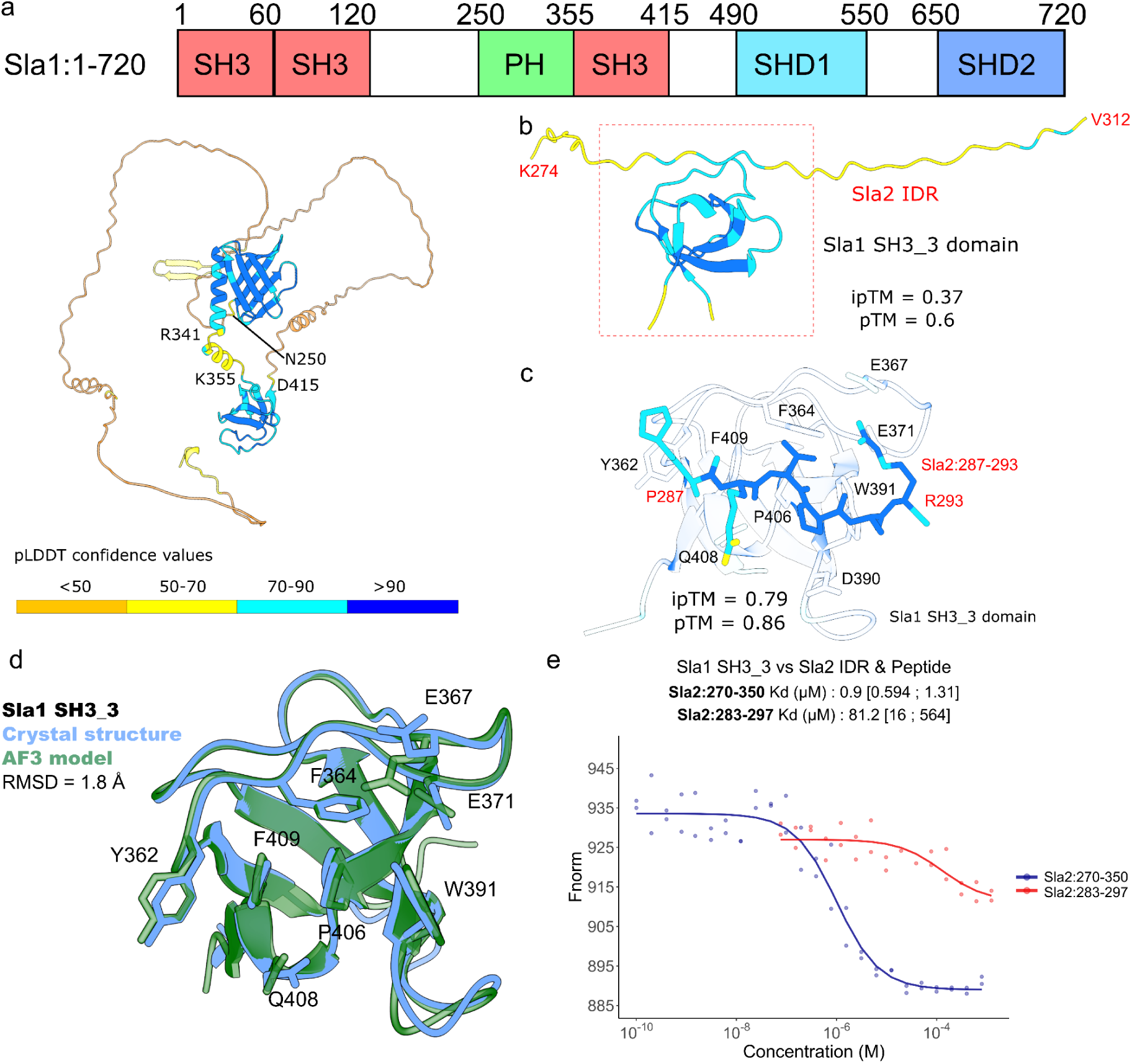
Sla1 SH3_3 domain binds the Sla2 IDR (A) Domain architecture of Sla1 residues 1-720 and AF3 model of Sla1 residues 120-510. (B) AF3 predicted complex of Sla1 SH3_3 and the extensive Proline rich IDR of Sla2 (residues 274-312). (**C**) The complex of Sla1 SH3_3 shown in grey and Sla2:287-293 was remodelled to reduce the error in the modelled interaction with the excess peptide region around the proline motif. The peptide is coloured as in the previous panels with the pLDDT score colour scheme. The contact residues between the motif and the SH3 domain are with the one-letter code and residue number. (**D**) Overlay of the crystal structure of the Sla1 SH3_3 domain (including a protein expression cleavage scar) with the AF3 model of the same amino acid sequence. The key residues that are found to be coordinating peptide motifs in other structures of SH3 domains are labelled and shown in stick form for both models. Aromatic residues spread through the binding cleft as well as two polar/charged residues that coordinate positively charged residues at the C-terminus of the Pro-rich motif. (**E**) MST of Sla2:270-350 and Sla2:283-297 titrated against Red-NHS labelled Sla1 SH3_3, which gave a binding K_d_ of 0.9 μM and 81.2 μM, respectively. These results confirm that the Sla2 IDR binds to the Sla1 SH3 domain.

Therefore, our hypothesis is that the primary interaction area from Sla2 would be the proline-rich disordered region between the ANTH and coiled-coil region of Sla2 forming a complex with the 3rd SH3 domain of Sla1. We modelled, using AF3, the residues 274 to 312 of Sla2 (the proline-rich region of the IDR) with residues 355 to 414 of Sla1 (SH3_3) (Figure 7b). The SH3_3 domain was modelled in proximity to residues 287-293 of Sla2, with an initially low iPTM score of 0.37. The iPTM score for the model of the Sla1 SH3_3 domain with the isolated peptide (residues 287 to 293) was 0.79 (Figure 7c). The key residues involved are similar to that of the crystal structure of an SH3 domain from *Hs*p40phox and a bound Proline rich peptide (*63*). The pTM score given by AF3 is a measure of confidence for the 3D position of all the atoms, whereas the iPTM score reflects the confidence for the positions of interacting atoms at an interface. A score above 0.6 for both, combined with additional experimental evidence, strongly indicates that the predicted model is likely to be correct.

We expressed and purified the SH3_3 domain of Sla1 (residues 355-414). Far UV Circular Dichroism spectrum and thermal denaturation experiments indicate that the domain is folded and displays a two state cooperative unfolding (*64*) (Supplementary Figure 9a-b). We solved the crystal structure of the Sla1 SH3_3 domain at 1.49 Å (Supplementary Figure 9c-e). The AF3 model and crystal structure have a high correlation as shown by the superimposed models in Figure 7d, with a RMSD between chain B of the crystal structure and the AF3 model of 1.8 Å. This well folded structure for the expression construct supports the confidence in our expression construct for use in biophysical analysis of the peptide interaction (Figure 7d). MST was used to investigate the interaction with the intrinsically disordered region (IDR) of Sla2, specifically the peptide 283-297 (Figure 7e). The K_d_ for the complete IDR (residues 270-350) is in the micromolar range, which is almost two orders of magnitude stronger. The change in affinity due to the inclusion of external regions from the peptide motif can be observed in other SH3 systems, such as with the GRB2 SH3 domains, where the complete IDR increases affinity for the peptide (*29*, *65*). Our biophysical evidence of the interaction, supporting crystal structure of the Sla1 SH3_3 domain, and AF3 models of the protein complex supports our proposal that the Sla1 SH3_3 domain interacts with the PARTPAR motif in the Sla2 IDR and that the complete IDR sequence has a positive impact on the SH3 domain Sla2 peptide interaction. As the Sla2 peptide N-terminus is the Proline residue coordinating with Y362 of the Sla1 SH3 domain and continues to the C-terminal Arginine residue that coordinates with E371, the PARTPAR motif is classified as a (-) direction binding motif of P^0^xxxP^4^xR^6^ (*27*). The motif PxxxPxR is sufficient for modelling the SH3 domain complex, with Alanine replacements of the non-proline residues showing little shift in the ipTM score apart from the terminal Arginine residue (Figure 8). In the IDR of Sla2/End4 proteins across Fungi this motif is present in 15 examples and a P^0^xxP^3^xR^5^ motif is more conserved with 155 examples in our selected sequences. The PxxPxR sequence is another common class of motifs bound by SH3 domains (*27*). In the IDR of Metazoa Hip1R sequences, only one example exists of the P^0^xxxP^4^xR^6^ and none of the P^0^xxP^3^xR^5^ motif within the IDR (Sequences used for alignment in Supplementary information).

**Figure 8:**
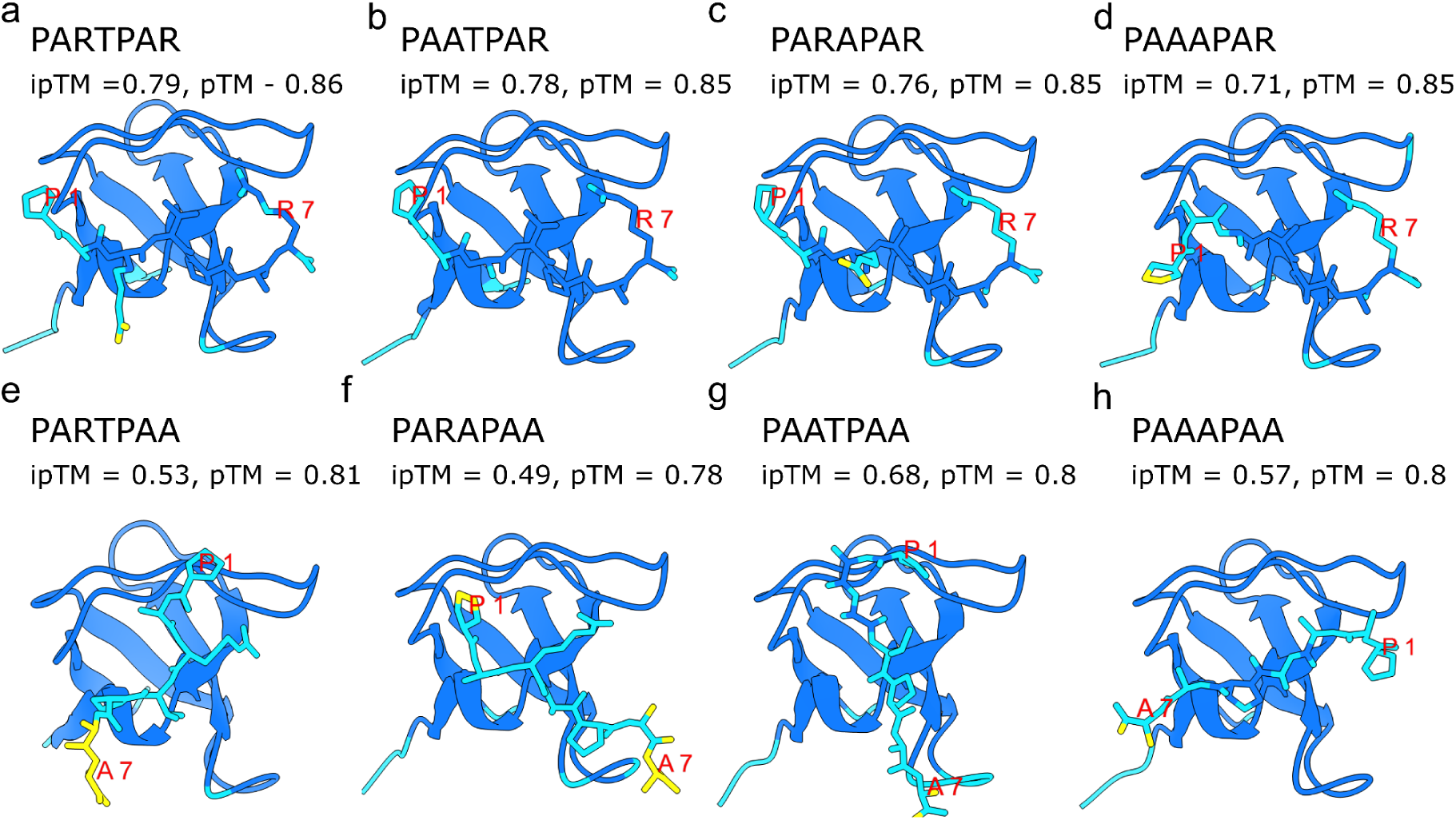
AF3 models highlight that the PxxxPxR motif found in Sla2 is sufficient to predict the Sla1 SH3_3:Sla2 complex (**A-H**) - AF3 models of Sla1 SH3_3 and seven residue peptides listed above the model along with the ipTM and pTM scores. Alanine replacements for residues in the motif show a decrease in the iPTM scores, particularly when the C-terminal Arginine is replaced with an Alanine. The colouring scheme is the AlphaFold3 pLDDT colour scale as used for the other models in this work.

The other player in the Pan1/End3/Sla1 complex that binds to Sla2 is Pan1. This protein is a very extended protein with many domains for interaction with adaptors and cargo in endocytosis (*59*, *66*). Previous work has shown that the Sla2:Pan1 interaction is mediated by the coiled-coils of both Sla2 and Pan1 (*26*). To investigate the strength of the interaction between Pan1 and Sla2, we purified Pan1:777-987 and titrated this construct against labelled Sla2cc for MST measurements (Figure 9a). Far UV Circular Dichroism of Pan1:777-987 and Sla2:270-350 indicates a predominant alpha helical content for Pan1:77-987 and confirms the intrinsically disordered nature of Sla2:270-350 (Supplementary Figure 9a). Our binding experiments showed that Sla2 has an affinity for Pan1 of 0.6 μM. The affinity is in line with parameters used in the Pan1 inhibition assay done by Toshima et al. where 100 nM of Sla2p was sufficient to partially inhibit Actin polymerisation by 50 nM of Pan1 in combination with 10 nM Arp2/3 (*26*).

**Figure 9:**
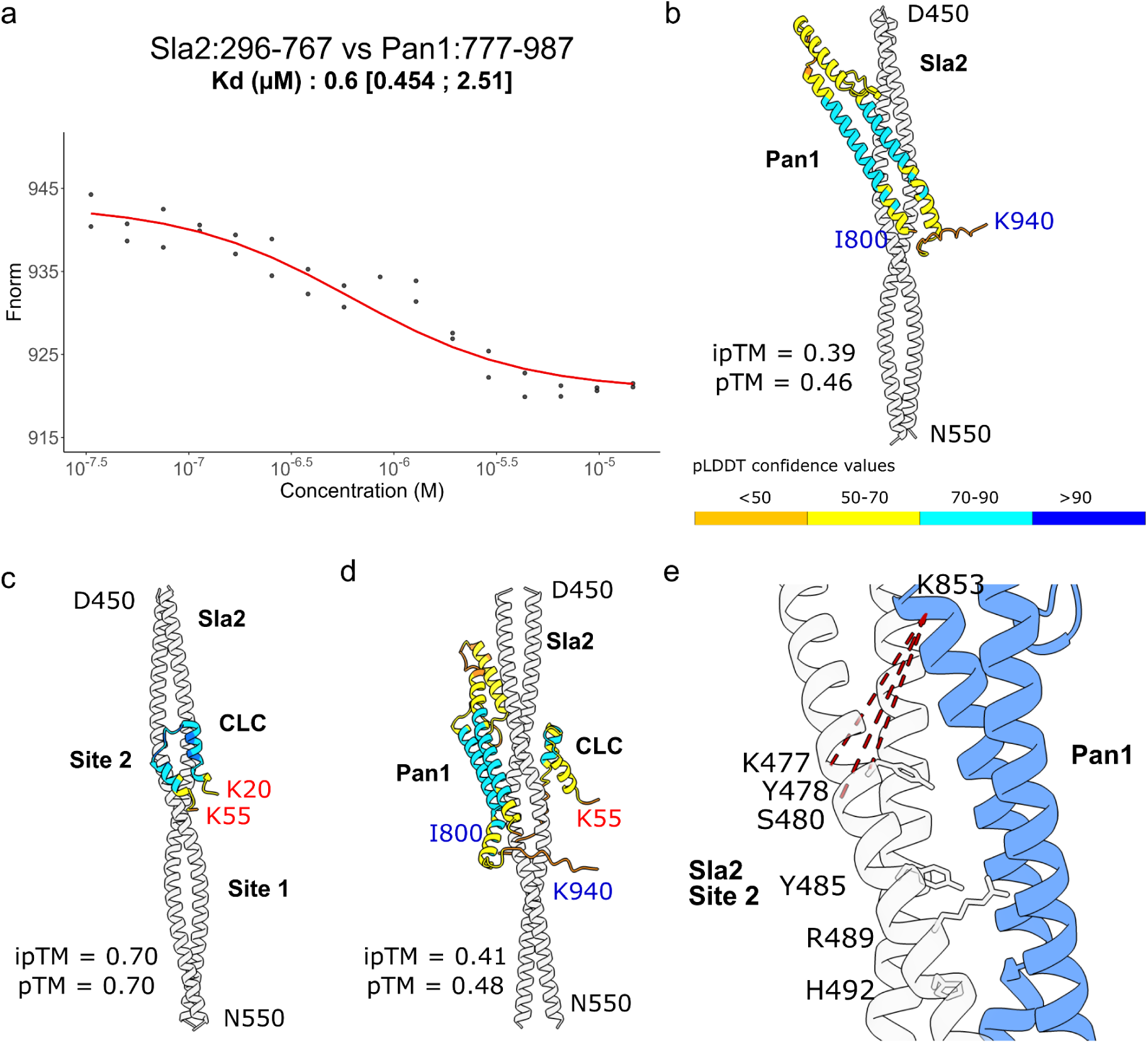
Pan1 and Sla2 interact through their coiled-coils, potentially at Site 2 (A) MicroScale Thermophoresis titration of Pan1:777-987 against RED-NHS labelled Sla2:296-767. This gave a resulting global K_d_ of 0.6 μM, CI95 [0.454; 2.51]. (**B-D**) AlphaFold3 models of Sla2:450-550, Pan1:800-940, and CLC combined in different ratios. (B) 2xSla2:CLC, (**C**) 2xSla2:Pan1, and (**D**) 2xSla2:CLC:Pan1. (**E**) Close up of the interface of the Sla2 dimer and Pan1 in (b). Key labelled residues in one Sla2 monomer are Y478, Y485, R489, and H492 that form Site 2 residues mutated in our two Site 2 mutant constructs, corroborating our hypothesis that Pan1 is competing for Site 2. Three cross-links between Sla2 and Pan1 are labelled in red dashed lines that were in the highest scored cross-links of our Pan1:Sla2 cross-linking dataset and correspond also to residues in proximity to each other in the AF3 model.

To complement the MST data, we used BS3 to cross-link Pan1 and Sla2 for Cross-Linking Mass-Spectrometry, which revealed a clustering of cross-links in the Site 2 region and at the C-terminal end of the construct in the first helix of the THATCH domain (Supplementary Figure 10). There were three times as many cross-links found in the Site 2 region as the Site 1 region. We subsequently modelled, using AF3, Sla2 residues 450-550, CLC residues 20-55 (Site 2 binding region of CLC), and Pan1 residues 800-940 in different ratios (Figure 9b-e). The complex of Pan1 with Sla2 is predicted with moderate-low confidence but with higher confidence in the individual folds of the polypeptides (Figure 9b). The models show that the CLC Site 2 complex is confident as the iPTM is above 0.6 in the Sla2:CLC complex (Figure 9c). Previous results from Skruzny et al., show by *in vivo* FRET measurements for the Sla2:Pan1 complex that the fluorophore labelled C-termini of both Sla2 and a truncated construct Pan1 (1–1050) have high FRET efficiency, showing that these protein termini are very close in physical space (*55*). This experiment suggests a suitable orientation to the model seen in Figure 9b. The reduced iPTM score of Sla2:Pan1 when compared to the Sla2:CLC complex may be the result of insufficient structural and sequence homology available in the AlphaFold3 model. The Pan1 moiety covers the same portion of Sla2 as CLC (Figure 9d). The location of the modelled interaction between Pan1 and Sla2 in both instances is predicted to be blocking Site 2, shown by the residues labelled on the Sla2 coiled-coil (Figure 9e). This is congruent with the high number of cross-links in this region between Pan1 and Sla2. In combination, these results show that Pan1 forms a coiled-coil that binds to Sla2 Site 2 with a comparable K_d_ to CLC.

## Discussion

Our results for the interaction of the Clathrin Light Chain to the coiled-coil of Sla2 is a diversion between Fungi and Metazoa. There are two independent binding sites of different magnitudes, the first that we characterised was the conserved site found in both Fungi and Metazoa.The second site that we found, Site 1, has a higher affinity than the conserved site and is directly C-terminal of Site 2 and binds the C-terminal region of CLC from Site 2. The conserved site (Site 2) has a clear role, the binding region in *Hs*CLC contains the EED (EQD in *Sc*CLC) motif that binds the knee of Clathrin Heavy Chain, which regulates the lattice stiffness (*40*, *43*, *44*, *67*, *68*). Our sequence alignment alongside the cross-links we found for the region of CLC and the AF3 model clearly show the acidic motif closely associated with the coiled-coil (Supplementary Figure 3). In contrast, the sequence alignment for Site 1 region in Sla2 and Hip1R (the human homolog of Sla2) shows little similarity (Figure 4a) and no high affinity complex was found by the prior work on Hip1R. In addition, the region for Site 1 in CLC is highly conserved between Fungi and Metazoa. Our results show that Site 1 is also the dominant of the two sites in yeast when mutated *in vivo*. The sequence similarity in CLC at Site 1 between Metazoa and Fungi, and not in Sla2/Hip1R may suggest that Site 1 emerged in Fungi due to an evolutionary pressure on competition for Site 2 that cannot lead to reciprocal changes in CLC as the interaction with the distal leg of CHC must remain unmodified.

We have solved the structure of the Sla2 C-terminal region. In the context of prior investigations into *Hs*Hip1R for Actin binding studies, the structure reveals that the THATCH domain Actin binding (ACB) surface is not freely available in the conformation we describe (Figure 6 and Supplementary Figure 6b). These residues form a cluster on the face of the THATCH core pointing in the direction of the coiled-coil (Figure 6b). As a result of the ACB facing the coiled-coil the Sla2 binding of Actin could act like a hook allowing the force of the cytoskeleton to be exerted and pull the membrane outwards similar to the model proposed for Talin (*47*). This helps relate the unfolding cycle of the REND domain and Up-Stream-Helix (USH) from the work of Ren et al. (*37*, *38*). The REND domain in our structure forms a dimer with a large interface between the two chains. This may be an influence on the large force required to unfold the domain, which may be additionally important when the LATCH helix has to unzip during the Actin binding process and the THATCH domains must remain close together during Actin binding. It is suggested that an increased local concentration of the REND domain has an activation effect as when it forms puncta on the membrane it forms potentially liquid-like droplets (*39*). The increased local concentration of Sla2 due to membrane recruitment may overcome the moderate affinity for Actin that was shown by Hip1R (*16*).

In the work of Ren et al., it was shown that the force redistribution in endocytosis can be mediated through the IDP and coiled-coil region of Sla2, from our data this would be the Pan1/End3/Sla1 complex, as Sla1 will bind to the IDR and Pan1 interacts with the fibrils (*38*). This changes the force on the REND and first helix, UpStream Helix (USH), of the THATCH domain and may affect the unfolding cycle on top of the force distributed by Actin on Sla2 (*23*, *26*). The force distributed by Sla1/End3/Pan1 interaction to F-Actin and subsequently Sla2 is roughly 8 pN, which is sufficient to unfold the USH. It is unclear whether that while CLC inhibits Sla2, this interaction modifies the force transmission to the USH. The unfolding of the USH would lead to an extended distance between the REND and THATCH domain allowing helix 3 and 4 to hook the Actin fibril. Taking the formula from Kohn et al. (*69*), L = 0.360x(Naa-1), we can surmise that the extended length of the unfolded linker between the REND and THATCH (with an unfolded USH) would be maximum 40 residues (residues 731-770), therefore L = 14 nm. The diameter of an Actin fibril is 7 nm, allowing for the Actin binding domains to hook around the fibres and be physically possible for the REND dimer to stay dimerised.The interaction of Pan1 has significant proximity to the CLC interaction with the coiled-coil and has a similar affinity. Pan1 can compete with Clathrin Light Chain for Sla2 in the endocytic pit. This could have been the selection pressure in Fungi that led to the co-evolution of CLC binding Site 1 in Sla2. Site 1 has a different physical location from Site 2 and a higher affinity for CLC, which relaxes the pressure on the competition for Pan1 at Site 2. Processive endocytic events are significantly decreased in the Sla2 Site 1 mutant as compared to the WildType, this could be explained by the effective competition for Site 2 increasing as only the Site 2 interaction remains for both proteins. Therefore, the Pan1 interaction may reduce the CLC occupation on Site 2 and lead to the regulatory pathway of Actin polymerisation being out of balance. An element of discussion about Site 1 is that the location in CLC is overlapping with the CLC central helix that binds to CHC. Whether the Clathrin heterodimer is weakened through this interaction or has some other allosteric effect on CHC would also be an open question because of our results.

The results we have presented here highlights Sla2 as an interaction hub with three key players of the endocytic coat. We have quantified the strength of these interactions between Sla2 and the respective binding partners as well as mapping these interactions to more specific regions than was previously possible. We can show a further understanding of the nature and strength of these interactions. Starting from the confirmed proline based motif in the IDR of Sla2 that binds the SH3_3 domain of Sla1 to enhance its role at the endocytic pit along with Pan1. Followed by the coiled-coil, which not only enables regulation of Actin binding through CLC interactions, it also interacts back to the Pan1/End3/Sla1 complex with Pan1. We find two interactions between Sla2 and CLC, one of which is conserved with Metazoa and a second higher affinity site that has no homology in Metazoa. Finally, we have also determined the complete structure of the C-terminal Actin binding region of Sla2 that puts the domains regulated by Clathrin Light Chain into three-dimensional context. Overall, these results provide additional insights in the complex network that dictates the outcome of endocytosis.

## Methods

### Protein production and purification

*Saccharomyces cerevisiae* cDNA was used for all protein expression apart from a codon optimised Sla1:355-414 cDNA, which was synthesised by GenScript (Piscataway, U.S.A.) into a pUC57 vector. Sla2cc constructs, Pan1:777-987, Sla1:355-414, and CLC constructs were expressed using pETM-30 containing a N-terminal 6xHis-Glutathione-S-Transferase-”TEV-cleavage-site”-tag. Sla2ccRTH was expressed using a pnEA N-terminal 6xHis-Glutathione-S-Transferase-”TEV-cleavage-site”-vector. NusA-CLC was expressed using a pnEA vector. Chemi-competent *E. coli* BL21 (DE3) cells already pre-transformed with the pLysS plasmid (Novagen) were transformed with 100 ng of plasmid DNA and grown overnight at 37 °C with 30 µg/ml of kanamycin (50 µg/ml ampicillin for the pnEA vector). For protein expression, a 1:100 dilution of the preculture was done in Terrific Broth medium (20 g tryptone, 24 g yeast extract, 4 ml Glycerol per litre, 0.072 M K_2_HPO_4_ and 0.017 M KH_2_PO_4_). Cells were grown at 37 °C until reaching OD 0.8. Then, the temperature was reduced to 16 °C, and induction was achieved by adding IPTG to a final concentration of 0.25 mM. Induction was performed overnight. Cells were harvested at 4,500 g for 20 minutes at 10 °C and stored at −20 °C until purification.

Cells were resuspended in 5 mL of lysis buffer (30 mM Tris pH 7.5, 300 mM NaCl, +400 U DNAse I, and a tablet of Complete EDTA-free protease inhibitor cocktail, Roche, per 100 mL of buffer) per gram of cells. Rupture of the cells was achieved by using an Emulsiflex C3 (Avestin, Ottawa, Canada) cell disruptor at 15 kPsi three times. A centrifugation at 40,000 g for 50 minutes at 4 °C was performed to clear the lysate. For purification of 6xHisNusA-CLC, the lysate was loaded onto a Ni-NTA (Carl Roth, Germany) gravity column equilibrated with buffer A (30 mM Tris pH 7.5, 300 mM NaCl, 5 % v/v glycerol, and 10 mM imidazole). The column was washed with 10 column volumes (CV) of buffer A and eluted with buffer B (30 mM Tris pH 7.5, 300 mM NaCl, 5 % v/v glycerol and 200 mM imidazole), collecting 1 mL fractions. The purity of the protein samples was assessed using SDS-PAGE. For Sla2cc, Sla2ccRTH, CLC (1-233, 1-80, 70-140, 70-233), Pan1:777-987, and Sla1:355-414 polypeptides, lysates were loaded onto GST-Sepharose4B gravity columns (Cytiva, Germany). Lysis buffer in these cases contains 30mM Tris pH 7.5, 300 mM NaCl, 5 % v/v glycerol, +400 U DNAse I and a tablet of Complete EDTA-free protease inhibitor cocktail, Roche, per 100 mL of buffer. Buffer A contains 30 mM Tris pH 7.5, 300 mM NaCl and 5 % v/v glycerol. On-column cleavage at 4 °C overnight was done after washing the column into 30 mM Tris pH 7.5, 300mM NaCl, 5 % v/v glycerol, 0.5 mM TCEP. For cleavage 1 mg of TEV per ml of beads was used.

The samples were loaded onto a Superdex 200 HiLoad 16/600 size-exclusion chromatography (SEC) column for *Sc*CLC and *Sc*Pan1 constructs; Sla2 constructs were loaded onto a Superose 6 HiLoad 16/600. Sla1 SH3_3 and Sla2 IDR were loaded onto a Superdex 75 10/300 size-exclusion chromatography column. In each case, the SEC column was equilibrated with SEC buffer (30 mM HEPES pH 8, 150 mM NaCl, 0.5 mM TCEP). Fractions were analysed for purity using SDS-PAGE, pooled, concentrated to between 5 and 20 mg/ml, flash frozen in liquid nitrogen, and stored at −70 °C until used.

Sla2:283-297 peptide was purchased from NovoProLabs (Shanghai, China) with a purity of at least 98% and TFA-free (Acetate-salt). Sequenced as PVSTPARTPARTPTP. The peptide was solubilized in SEC buffer at 10 mM concentrations.

### MicroScale Thermophoresis

CLC, Sla1:355-414 and Sla2cc were labelled using the 2nd Generation RED-NHS labelling kit (Nanotemper, Munich, Germany). The reaction was done in SEC buffer, in the dark, at room temperature, for 30 minutes, with a 3-fold excess of dye used per molecule of protein. Proteins were passed through the enclosed PD-10 columns provided by the manufacturer to remove unlinked dye from the protein samples and to restore the proteins back into SEC buffer. Proteins were directly aliquoted from the elution fractions and frozen in liquid nitrogen for −70 °C storage. Concentrations between 25-100 nM of each labelled protein, depending on the label efficiency for each polypeptide, were used for titrations against ligands at 25 °C. Samples were incubated together for 5 minutes prior to measurement. In the case of labelled CLC the titrations were against Sla2cc (WildType, Site 2 Mutant 1, Site 2 Mutant 2, deltaSite1). For labelled Sla2cc it was CLC:70-140, CLC:70-233, and Pan1:777-987. Labelled Sla1:355-414 was titrated against unlabelled Sla2:270-350 and Sla2:283-297. MST data was exported and compressed into *zip file* for each set of analyte and ligand MST runs (n=2) and loaded into ThermoAffinity (spc.embl-hamburg.de) for data analysis before exporting the fitting data and replotting in R (*70*).

### Thermal Stability assay / nanoDifferential Scanning Fluorimetry (nDSF)

Sla1:355-414 was diluted to 20 μM in the SEC buffer described in the purification protocol. 3 capillaries with 10 μL each were inserted into the Prometheus nanoDifferential Scanning Fluorimeter (NanoTemper, Munich, Germany). A temperature ramp from 20-95 °C was performed, the temperature range 20 to 85 °C was used for analysis due to noise at the high temperature range. Processed data was exported from the Prometheus software and loaded into MoltenProt (*70*, *71*) for Tm fitting. Processed 350 nm signal data was replotted in R, averaging the three replicates for 350 nm signal.

### Mass Photometry

Constructs of Sla2:296-767, 6xHisNusA-CLC, and 6xHis-NusA were measured separately and in combination using a 1:19 dilution into the SEC buffer from a stock solution of 1 μM. The proteins were all purified into the final SEC buffer as in the protein purification section. Refeyn 2.0 (Refeyn, Oxford, United Kingdom) was used to collect. After loading the clean coverslip, 19 μL of buffer is applied to the sample compartment for focusing before mixing 1 μL of sample. Mass-contrast calibration was done prior to data export using Native protein marker (Thermofisher). Data analysis was performed on the calibrated movies using the eSPC tool Photomol, where gaussian fitting and binning of the data was performed before replotting in R (*51*).

### Circular Dichroism

Sla2cc constructs, Sla1:355-414, Sla2:270-350, and *Sc*Pan1:777-987 were dialysed overnight into 30 mM NaPO_4_ pH 7.5, 150 mM NaF using a Slide-A-Lyzer MINI Dialysis Device 10K MWCO 0.5 ml (Thermo Scientific). These samples were then measured from 180-300 nm in a 400 μL, 1 mm pathlength quartz cuvette (Hellma, Müllheim, Germany) using a ChiraScan Circular Dichroism Spectrophotometer (Applied Photophysics, Leatherhead,UK), equipped with a Quantum Northwest TC 125 temperature controller (Liberty Lake, Washington, USA) set at 20 °C. Sla2cc WildType and Sla2:270-350 were measured at 0.25 mg/ml and all other samples were measured at 0.125 mg/ml. Three replicates of the spectrum were recorded, each with a step size of 1 nm and a response time of 1 second. Temperature ramps from 20-90 °C for Sla2cc constructs, 1 °C per minute increase and CD measured at 5 °C intervals. All data was buffer-subtracted and converted to Mean Residue Weight Extinction for analysis to be able to compare samples of different concentrations and residue length. The ChiraKit server from the eSPC online toolkit was used to analyse the data for Melting Temperature and Secondary Structure contents https://spc.embl-hamburg.de/app/chirakit.

### Generation of mutations of pETM-30-Sla2cc and pRS315-5’UTR Sla2-mScarlet-I

Mutagenesis was performed with the following criteria for Site 2 Mutant 1: all residues were to be mutated to alanines. For Site 2 Mutant 2 the replacements were designed to switch the charge of the side chain of charged residues and replace polar and aliphatic residues with an alanine. With the one exception of R498 due to sequence restrictions, which was replaced by a glycine. Mutants were generated by Quikchange mutagenesis. Whole plasmid PCR was performed using overlapping primers, subsequently 0.5 μL of Dpn1 restriction enzyme was added to digest the original vector. *E. coli* cells were transformed with the PCR mix and selected on LB agar plates before subsequent picking and sequencing by Sanger sequencing (Microsynth, Göttingen, Germany.

### Cross-linking and LC-MS/MS

Sla2cc+CLC:1-80 and Sla2cc+Pan1:777-987 cross-link samples were mixed in equimolar amounts to a final 20 μM total protein concentration. BS3 X-linker was added at 0.5 mM and incubated for 30 mins at 25 °C. Quenched with 100 mM Tris pH 8 for an hour. Samples were then sent at room temperature to the Proteomics Core Facility in Heidelberg for desalting and Peptide-Size exclusion chromatography.

For the digestion, 5 mM TCEP, 20 mM CAA and 1 µg trypsin were added and incubated at 37 °C overnight. Next day, reaction was stopped by the addition of 1 % TFA. Digested peptides were concentrated and desalted using an OASIS® HLB µElution Plate (Waters) according to manufacturer instructions. Crosslinked peptides were enriched using size exclusion chromatography(*72*). In brief, desalted peptides were reconstituted with SEC buffer (30 % (v/v) ACN in 0.1 % (v/v) TFA) and fractionated using a Superdex Peptide PC 3.2/30 column (GE) on a 1200 Infinity HPLC system (Agilent) at a flow rate of 0.05 ml/min. Fractions eluting between 50-70 μL were evaporated to dryness and reconstituted in 30 μl 4% (v/v) ACN in 1 % (v/v) FA.

Collected fractions were analysed by liquid chromatography (LC) - coupled tandem mass spectrometry (MS/MS) using an UltiMate 3000 RSLC nano LC system (Dionex) fitted with a trapping cartridge (µ-Precolumn C18 PepMap 100, 5 µm, 300 µm i.d. x 5 mm, 100 Å) and an analytical column (nanoEase™ M/Z HSS T3 column 75 µm x 250 mm C18, 1.8 µm, 100 Å, Waters). Trapping was carried out with a constant flow of trapping solvent (0.05 % trifluoroacetic acid in water) at 30 µL/min onto the trapping column for 6 minutes. Subsequently, peptides were eluted and separated on the analytical column using a gradient composed of Solvent A ((3 % DMSO, 0.1 % formic acid in water) and solvent B (3 % DMSO, 0.1 % formic acid in acetonitrile) with a constant flow of 0.3 µL/min. The outlet of the analytical column was coupled directly to an Orbitrap Fusion Lumos (Thermo Scientific, SanJose) mass spectrometer using the nanoFlex source. The peptides were introduced into the Orbitrap Fusion Lumos via a Pico-Tip Emitter 360 µm OD x 20 µm ID; 10 µm tip (CoAnn Technologies) and an applied spray voltage of 2.1 kV, instrument was operated in positive mode. The capillary temperature was set at 275 °C. Only charge states of 4-8 were included. The dynamic exclusion was set to 30 sec. and the intensity threshold was 5e^4^. Full mass scans were acquired for a mass range 350-1700 m/z in profile mode in the orbitrap with resolution of 120000. The AGC target was set to Standard and the injection time mode was set to Auto. The instrument was operated in data dependent acquisition (DDA) mode with a cycle time of 3 sec between master scans and MSMS scans were acquired in the Orbitrap with a resolution of 30000, with a fill time of up to 100 ms and a limitation of 2e5 ions (AGC target). A normalised collision energy of 32 was applied. MS2 data was acquired in profile mode.

All data was analysed using the cross-linking module in Mass Spec Studio v2.4.0.3524 (www.msstudio.ca, doi: 10.1074/mcp.O116.058685). Parameters were set as follows: Trypsin (K/R only), charge states 4−8, peptide length 7−50, percent Evalue threshold = 50, MS mass tolerance = 10 ppm, MS/MS mass tolerance = 10, elution width = 0.5 min. BS3 cross-links residue pairs were constrained to KSTY on both sides. Identifications were manually validated, and cross-links with an E-value corresponding to <0.05% FDR were rejected. The data export from the Studio was filtered to retain only cross-links with a unique pair of peptide sequences and a unique set of potential residue sites.

### Grid preparation

Sla2ccRTH was diluted into SEC buffer at a concentration of 20 μM. For cryo-EM grid preparation, Quantifoil 300 mesh Au R 2/2 holey carbon grids were glow-discharged in a Cressington 208 carbon coater at 10 mA and 0.1 mbar air pressure for 60 s. The sample was then applied to the grid and vitrified using a Vitrobot mark IV (FEI/Thermo Scientific) with a blot force of −3 and a blot time of 3 s. The relative humidity (RH) was ≥ 90 % and temperature 5–6 °C. Liquid ethane was used as the cryogen.

### Electron cryo-microscopy and data processing

A Krios G3i electron microscope (FEI/Thermo Scientific) at the Centre for Structural Systems Biology (CSSB) Cryo-EM facility, operated at an accelerating voltage of 300 keV equipped with a K3 BioQuantum (Gatan) filter using a Falcon III direct electron detector operating in integrating mode for data collection. Cryo-EM data were acquired using EPU software (Thermo Fisher) at a nominal magnification of 120,000 X, with a pixel size of 0.68 Å per pixel. Movies of a total fluence of ∼ 45 electrons per Å^2^ were collected at a dose of 1 e^-^/Å^2^ per frame. A total number of 5,725 movies for the dataset was acquired at an underfocus range of 0.5 to 2.0 μm.

Processing of the Sla2ccRTH data was done using cryoSPARC v4 (*73*). All final parameters for the model are found in (Supplementary Table 1). The general pipeline followed several stages. Micrographs were corrected for beam-induced motion using patch motion correction and patch CTF correction in cryoSPARC. Particles were picked initially using the blob picker at 120-180 Å in diameter and particles were extracted using 1.5 times the size of the blob diameter. To generate 2D classes and a 3D reconstruction, several rounds of 2D classification and selection was done. Then the best 2D classes were used to do template picking on the micrographs. A subset of these was used to generate an initial model and all particles were subjected to rounds of 3D classification to remove ‘bad’ particles via heterogeneous 3D refinement tools. Around 250,000 particles were taken forward to model refinement in C2 symmetry. Non-uniform refinement in cryoSPARC v4 was used to further improve the resolution of the density map and the final reconstruction was sharpened with a Resolution determined filter after the final Local Refinement job. Model building into the density map given from this dataset was done using an initial AlphaFold3 model for residues 560-968 of Sla2 (Supplementary Data) and manipulation in ChimeraX with the Molecular Dynamics tool ISOLDE(*74*). Refinement of the model was completed in PHENIX (*75*). The model was uploaded with the relevant density maps into the wwPDB for resolution estimation and final reports for model fitting.

### Computational modelling and alignment of Sla2, CLC, Sla1, and Pan1

Jalview was used to visualise sequence alignments of CLC and Sla2 sequences and homologous protein sequences using MAFFT and MUSCLE alignment parameters (*76*, *77*). Sequences for alignment were provided using UniprotKB; for CLC 100 representative sequences were found for both Fungi and Metazoa after screening sequences for maximum 95 % identity and length between 200-400 residues. For Sla2, Sla2/End4 protein names were used to find Fungal sequences in UniProtKB and Hip1R in Metazoa below 95 % identity and length between 800-1200 residues. The residues were regrouped after alignment of all the sequences back into Fungal and Metazoa and coloured in each group based on conservation. AlphaFold3 (AF3) online servers were used for the structural models. FoldSeek was used on AF3 models of Sla1 residues 251-360 and 2nd OPY1 PH domain for bidirectional search within the yeast structural proteome (*60*).

### Protein crystallisation and model refinement

Sla1 SH3_3 was crystallised in 0.1 M HEPES pH 7.5 and 20% PEG 10000. Crystals formed overnight at 19 °C and harvested after 3 days. The dataset was collected at P13 operated by EMBL at the PETRA III storage ring (DESY, Hamburg, Germany) (see Supplementary Table 3). The dataset was reduced and scaled using AIMLESS(*78*). The structure was solved using molecular replacement, using MOLREP(*79*) with the AF3 model of Sla1_SH3_3 expression construct with the GAMA cleavage scar as the search model. Iterative refinement and model-building cycles were performed using REFMAC(*57*) and Coot(*80*) in the CCP4i2 suite of programs(*81*).

### Measurement of endocytic dynamics

#### Generation of competent yeast cells with endogenously tagged Ent1p-mNeonGreen and Abp1-mTurquoise2

A 5 mL preculture of cells was incubated overnight at 30 °C with shaking in the appropriate medium. The OD of the overnight culture was determined and a new 50 ml culture with fresh medium was inoculated to OD of 0.15. Cells were grown at 30 °C until reaching an OD_600nm_ between 0.4 and 0.6. Cells were spun down at 3000 g, at 4 °C for 5 minutes, discarding the supernatant. Cells were resuspended in 30 mL of sterile water and spun down again at 3000x g, at 4 °C for 5 minutes, and the supernatant was discarded. Cells were resuspended in 1 ml of sterile water and spun down in a tabletop centrifuge at 4 °C for 5 minutes at 3000 g, and the supernatant was discarded. Cells were resuspended in 300 μL competent cell solution (5 % v/v glycerol; 10 % v/v DMSO), aliquoted in 50 μL, and stored at −70 °C (cooled down as slowly as possible).

#### Generation of sla2pΔ cells expressing pRS315-Sla2_mScarlet

Competent *Saccharomyces cerevisiae* cells (MK100 WT: MATa; his3Δ200; leu2-3,112; ura3-52; lys2-801) with Ent1p endogenously tagged with mNeonGreen and Abp1 tagged with mTurquoise2 were transformed with a vector containing the 5’ UTR of Sla2, the ORF of Sla2 fused in the 3’ with mScarlet (pRS315-Sla2_mScarlet) containing the LEU auxotrophy factor cassette using the following protocol (*82*). Cells were grown overnight in YPD media and fresh YPD media was inoculated to an OD 600 nm of 0.1-0.15. The culture was grown until an OD600nm of 0.4-0.6 was reached. Cells were spun down at 3000 g for 5 min, the supernatant was discarded, the pellet was resuspended in 30 mL cold H2O and spun down as before described, supernatant was discarded. Cells were resuspended with 1 mL of 100 mM lithium acetate and transferred to a 1.5 mL centrifuge tube, spun down for 15 s in a tabletop microcentrifuge. Supernatant was discarded and 400 μL of 100 mM lithium acetate was added to resuspend the cells. 50 μL of the cell suspension was mixed with precooled 240 μL 50 % PEG 3350, 10 μL of plasmid pRS315-Sla2_mScarlet carrying Sla2 gene tagged with mScarlet and LEU selection marker and 25 μL of 2 mg/mL ssDNA was added and vortexed. Cells were incubated at 30 °C for 30 min and at 42 °C for 25 min. Transformation solution was spun down for 15 s, supernatant was discarded and 100 μL H2O were added to plate cells on SC-Leu Agar plates. The same protocol was followed for transformation to knock out endogenous Sla2, only the cells before transformation were grown in SC-Leu medium. Once colonies were found, competent cells were prepared and transformed with a URA resistance cassette with homology arms to endogenous Sla2 gene (See Supplementary Table 2 for primer S1/S2 sequence) using the established PCR cassette protocol (pFA6a-KlURA3 Vector)(*83*).

#### Lifetime TIRF microscopy

To determine the effect of mutating the Sla2 coiled-coil in how endocytosis progresses, cells tagged with Ent1p-mNeonGreen, Sla2-mScarlet and Abp1-mTurquoise2 were used. Yeast cells previously described were grown overnight at 30 °C with shaking in a 24-well plate using LD(=low-fluorescence SD)-Trp-, Leu-medium (yeast nitrogen base without amino acids supplemented with the corresponding DropOut media, Foredium, CYN402). Cells were diluted in fresh medium with a starting OD600 nm of 0.1 and allowed to grow at 30 °C with shaking for several hours (4-6hs) until they reached log phase (OD600 nm 0.6-1.2). Micro slide 8-well glass bottom plates (Catalogue 80807, Ibidi, Gräfelfing, Germany) were treated with 50 μL of a 1 mg/ml Concanavalin A (prepared in 10 mM sodium phosphate buffer pH 6, 10 mM CaCl2,1 mM MnCl2, 0.01 % NaN3, Catalog C2010, Sigma Aldrich) solution, incubated for 5 minutes and then washed twice with 50 μL of fresh medium. 50 μL of cell suspension was applied, incubated for 5 minutes and removed. Then each well was washed twice with 50 μL of fresh medium. Finally, 50 μL of fresh medium was added.

TIRF Microscopy was done at room temperature (21 °C) using a Nikon Eclipse Ti2 microscope equipped with 405 nm and 488 nm lasers and an ORCA-Fusion BT Digital CMOS camera installed in the Advanced Light and Fluorescent Microscopy (ALFM) Facility in CSSB Hamburg (DESY, Hamburg, Germany). An oil immersion 100x objective was used (NA 1.49). For each field of view, a 5-minute movie was taken. Exposure for each channel was 500 ms in a 2 s interval (0.5 fps for each channel). Depending on cell density, movies were taken in 7 to 10 fields of view. Background subtraction was done with FIJI68. cmeAnalysis(*56*) was used to track and classify the tracked particles., These events were further classified into Abp1 positive and Abp1 negative events using the dual colour tracking functionality available in the package. Running parameters for tracking and classification were kept as default by the package. Data was plotted with ggplot2 and Pairwise comparisons were done using a Mann–Whitney U-test implemented in R(*84*).

## Acknowledgments

We thank Prof. Dr. Henning Tidow for suggestions and manuscript proofreading. We acknowledge the staff of the Sample Preparation and Characterisation (SPC) facility of EMBL Hamburg at PETRA III (DESY, Hamburg) for assistance (Stephan Niebling, Angelica Struve, Osvaldo Burastero, and Katharina Veith). We thank the EMBL Proteomics Core Facility (European Molecular Biology Laboratory), especially M. Rettel, for data acquisition and analysis. The electron cryo-microscopy was performed at the Multi-User CryoEM Facility at the Centre for Structural Systems Biology, Hamburg, supported by the Universität Hamburg and DFG grant numbers (INST 152/772-1|152/774-1|152/775-1|152/776-1|152/777-1 FUGG), the Federal Ministry of Education and Research (BMBF) and the DLR Projektträger (project SEEK 01KX2220). We acknowledge the support, technical assistance and advice of the CryoEM Facility staff. We thank the staff of the EMBL P13/14 beamline. The synchrotron data was collected at the P13 beamline operated by EMBL Hamburg at the PETRA III storage ring (DESY, Hamburg, Germany).

## Author Contributions

G.D.B produced the proteins for Mass Photometry, Microscale Thermophoresis, Nano Differential Scale Fluorimetry, Circular Dichroism, and Cross-Linking Mass Spectrometry. G.D.B. performed biophysical and structural experiments, and interpreted X-ray crystallographic & cryo-EM data. E.G. reviewed the cryo-EM data and model curation under the supervision of M.L.. L.A.D performed Total Internal Reflection Fluorescence microscopy and D.R.C. reviewed crystallographic data collection and model refinement. G.D.B. and M.G.A. conceived the project. M.G.A. supervised the project. G.D.B. and M.G.A. wrote the manuscript with input from all the authors.

## Data and material availability

All data supporting the findings of this study are available within the paper and its supplementary information. X-ray crystallographic data has been deposited in the PDB with code 9HDB and cryo-EM data is likewise deposited in the PDB and EMDB with code.

