## Supplementary information for "Sla2 is a core interaction hub for Clathrin Light Chain and the Pan1/End3/Sla1 Complex"

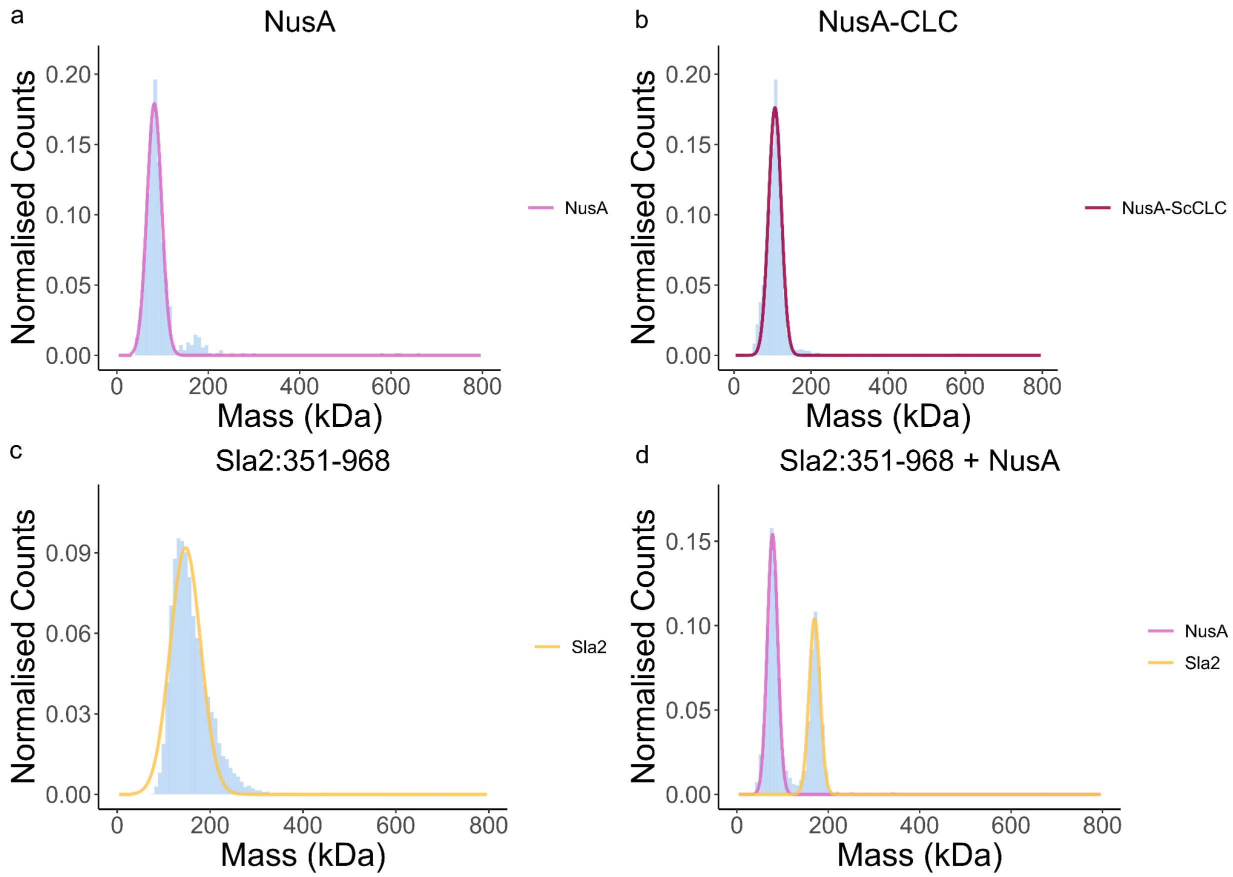


**Supplementary Figure 1: Controls for Mass Photometry visualisation of Sla2:CLC complex**

(**A**) NusA is a monomer at 82 kDa (σ = 16 kDa). (**B**) NusA-CLC is monomeric as well with a monomeric mass of 106 kDa (σ = 17 kDa). (**C**) Sla2:351-968 is dimeric with a mass of 147 kDa (σ = 33 kDa). (**D**) The NusA moiety does not bind to the coiled-coil. NusA was measured at 78 kDa (σ = 11 kDa) and Sla2 at 170 (σ = 12 kDa). The mass of the expected NusA moiety is 56 kDa, NusA-CLC is 85 kDa, and Sla2:351-968 dimer is 138 kDa. Gaussian fitting achieved through the eSPC program, PhotoMol.

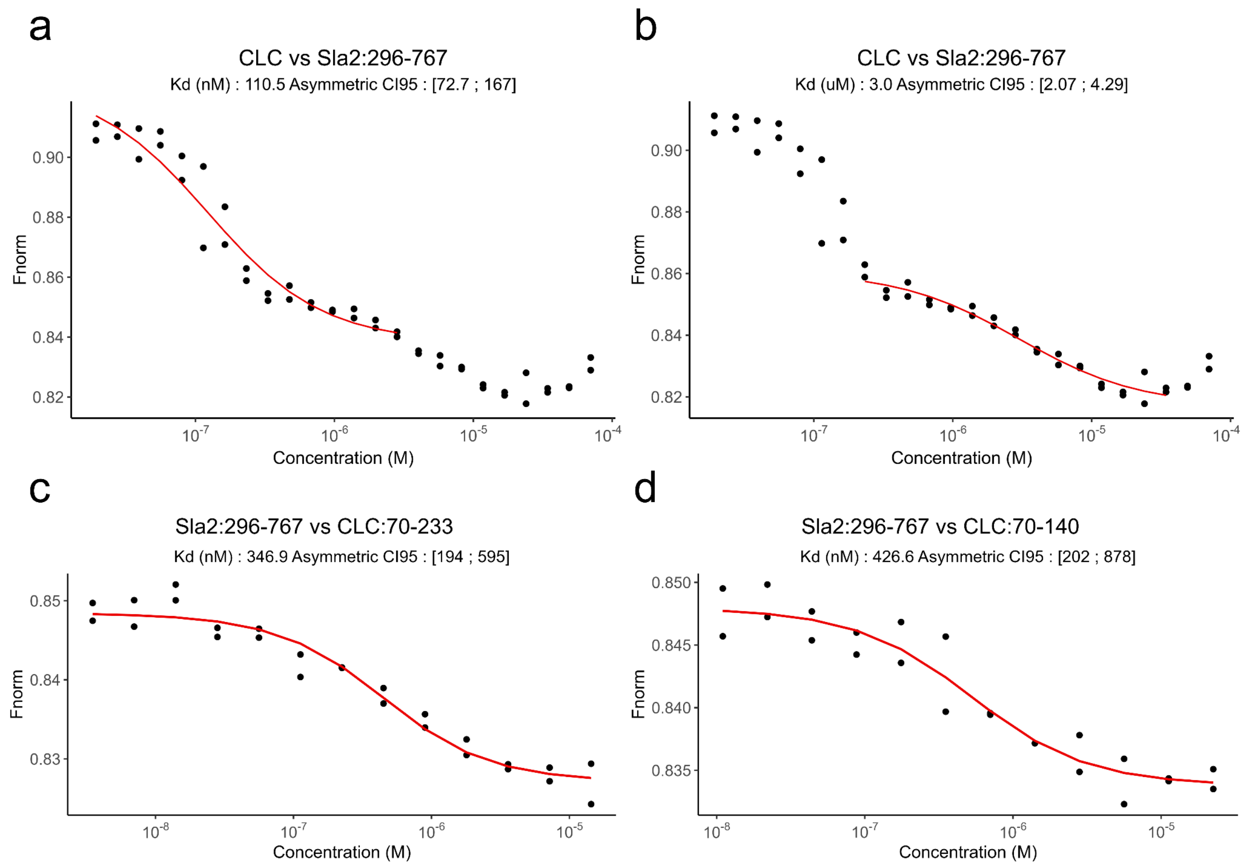


**Supplementary Figure 2: Microscale Thermophoresis maps the locations of two binding sites for Clathrin Light Chain:**

(**A**-**B**) Sla2cc titrated against Clathrin Light Chain-Red-NHS labelled for MicroScale Thermophoresis. Two transitions were measured and separately determined in the hundred nanomolar range and low micromolar range. (**C**) CLC residues 70-233 titrated against Sla2cc-REDHS. (**D**) CLC:70-140 titrated against Sla2cc-REDNHS. These gave similar sub-micromolar $K_d$s. These results show that CLC and Sla2 have two interaction interfaces, located in residues 1-70 and 70-140 of CLC. The measured affinities for these sites are an order of magnitude in difference; ~100-200 nM for Site 1 and 3.0 μM for Site 2.

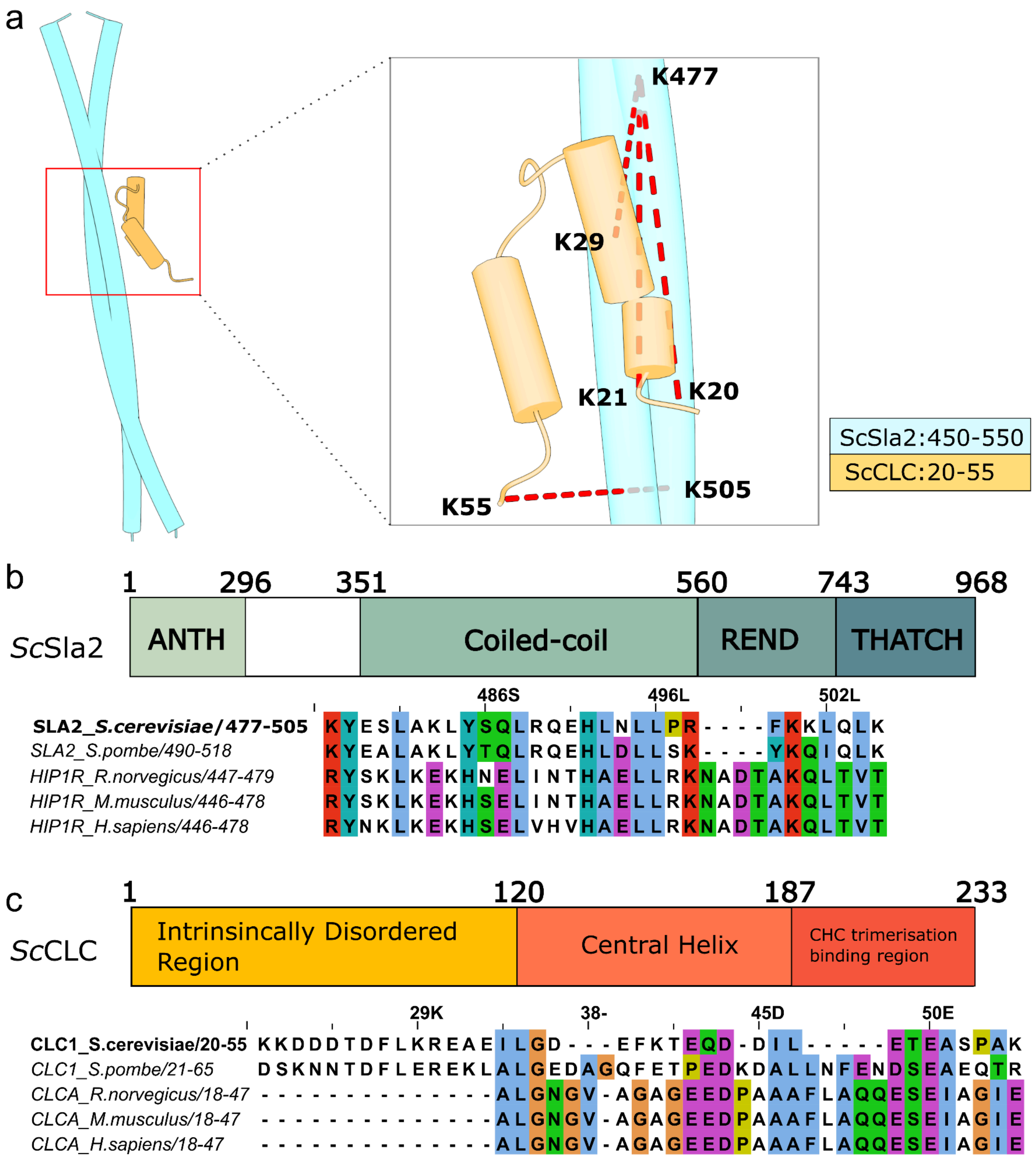


**Supplementary Figure 3: Site 2 is conserved in Sla2/Hip1R and CLC across yeast and mammals**

(**A**) AlphaFold 3 predictions of the dimeric Sla2 coiled-coil (450-550) and monomers of CLC were compiled and chemical cross-linking data is mapped onto the model. The high confidence inter-molecular hits from BS3 cross-linking experiments are subsequently labelled (red dashed lines). (**B**) Sla2 domain architecture is labelled with residue numbering alongside a sequence alignment of the cross-linked region of Sla2 from the Sla2:CLC BS3 cross-linking experiments with two model yeast sequences and three mammalian model organisms shown in the figure.

(**C**) The CLC architecture is labelled alongside the alignment of the cross-linked area of residues 20-55 with the same model organism protein sequences aligned to the *Saccharomyces cerevisiae* sequence.

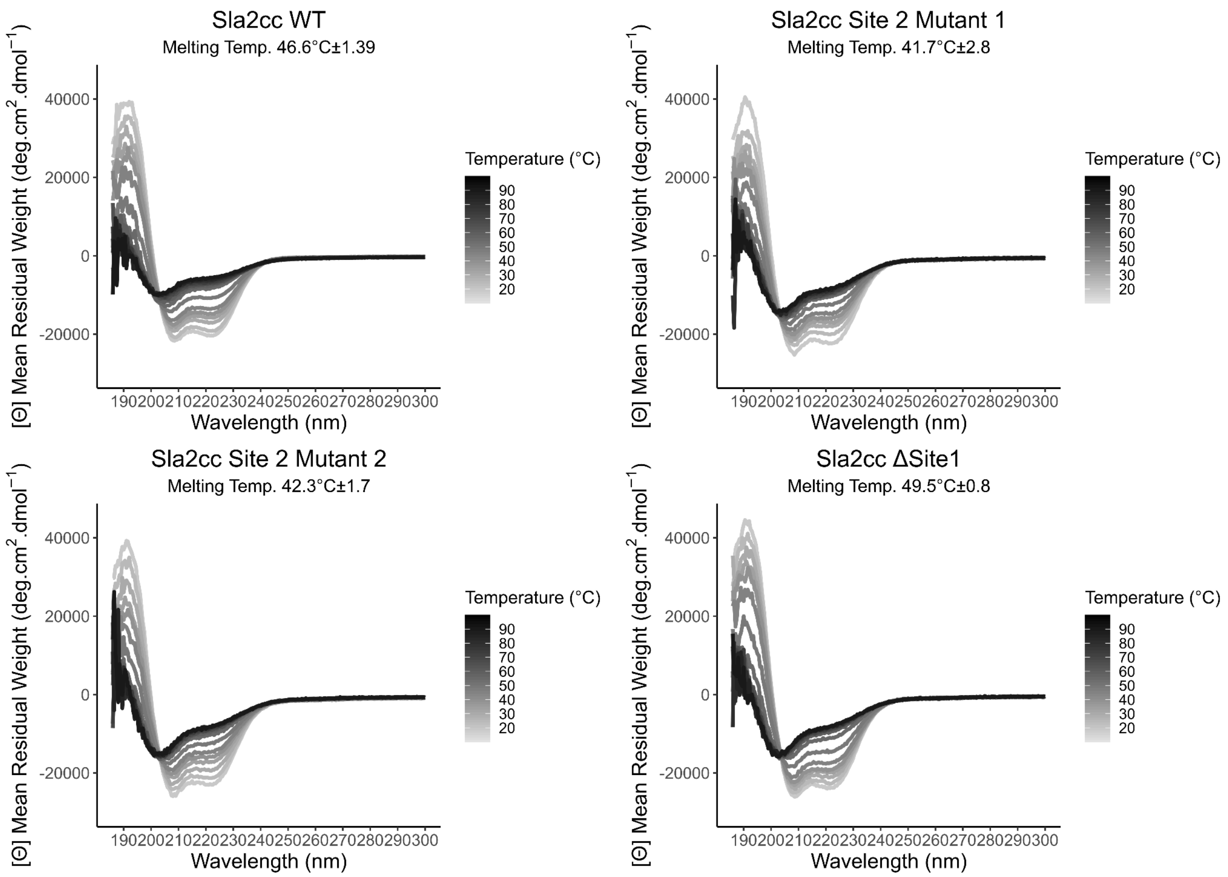


**Supplementary Figure 4: Circular Dichroism of Sla2cc for comparison of WildType to mutant constructs used for biophysical characterisation**

Circular Dichroism temperature ramps from wavelengths 180 nm to 300 nm. (**A**) Sla2cc WT, (**B**) Sla2cc Site 2 Mutant 1, (**C**) Sla2cc Site 2 Mutant 2, (**D**) Sla2cc ΔSite1. Circular Dichroism was performed to determine if the mutants used to characterise the CLC binding sites were folded correctly. Due to background noise, the spectra were used from 186 nm onwards. The constructs were all folded in similar secondary structure contents at 20 °C, with between 52 % and 58 % alpha helical content, the rest was distributed primarily between disordered regions and some low amounts of beta sheets and turns. The Far UV CD curves between 20 °C and 90 °C provided a suitable basis for fitting a melting temperature to each construct. The stabilities do not drop below 40 °C, but the Site 2 mutants do drop as compared to the WildType, and the Site 1 mutant increases the Sla2cc melting temperature by 3 °C. Fittings for melting temperature and secondary structure prediction from the Circular Dichroism curves were done using ChiraKit from the eSPC online toolkit https://spc.embl-hamburg.de/app/chirakit.

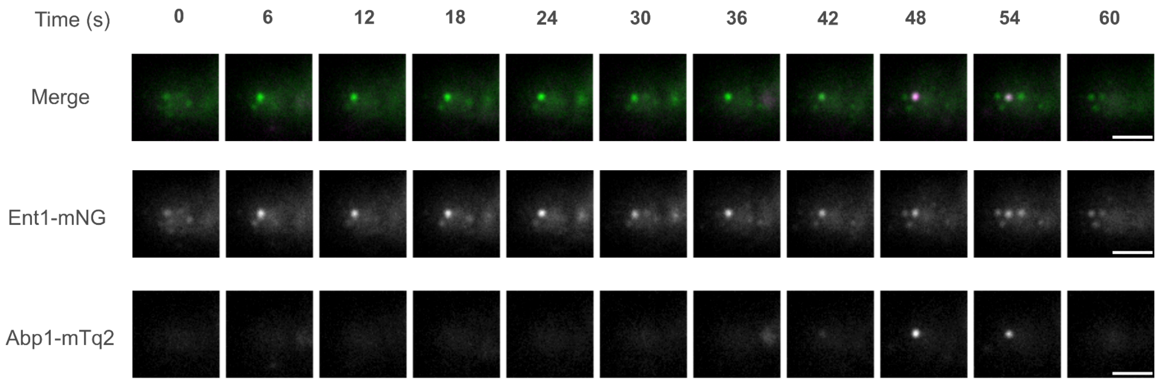


**Supplementary Figure 5: TIRF microscopy of endocytic events using Ent1-mNeonGreen and Abp1-mTurqouise2 to classify productive events**

A time-lapse sequence capturing a single endocytic event that demonstrates the co-localization of Ent1 fused with mNeonGreen (displayed in green) and Abp1 fused with mTurquoise2 (shown in magenta) in a wild-type Sla2 cell.

a

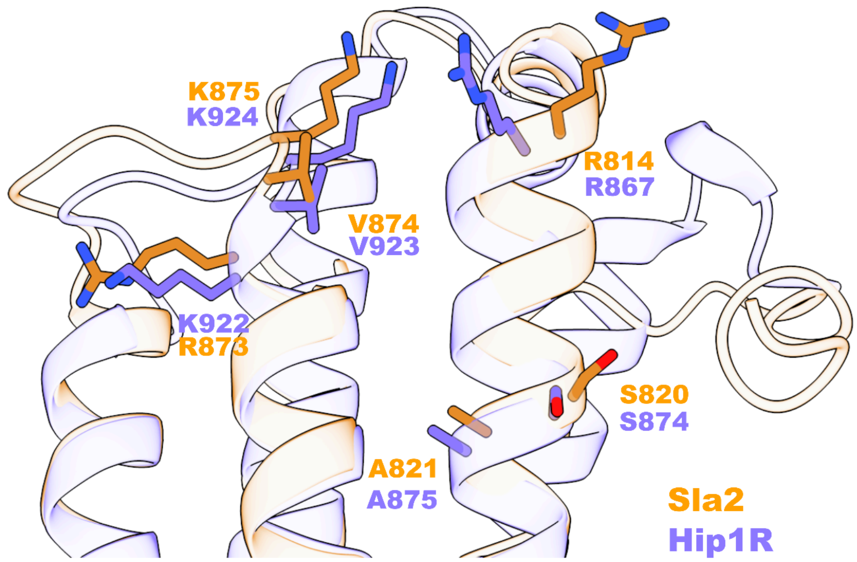


b

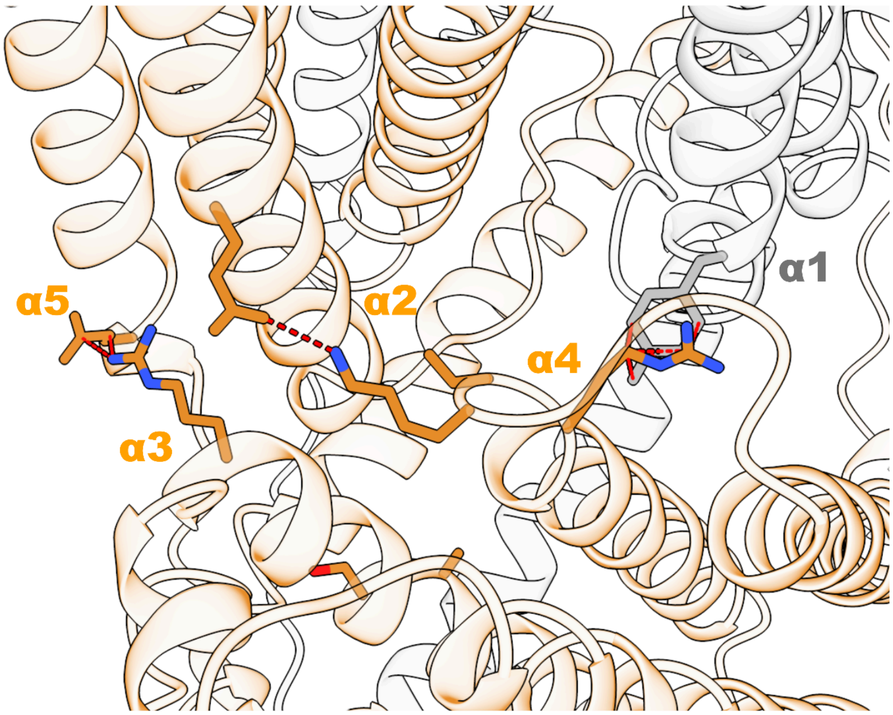


**Supplementary Figure 6: The THATCH domain Actin binding surface contacts the REND domains of both chains**

(**A**) Overlay of the THATCH domain structures of Sla2 and Hip1R, with Actin binding residues highlighted and labelled. The Sla2 THATCH domain is orange and the Hip1R crystal structure is purple. The positions have an RMSD of 0.8 Å. (**B**) The Sla2 REND and THATCH domains shown with Actin binding residues and their contacts to the REND domains highlighted with red dashed lines. The context of the other domains in our structure show that the ACB of one chain contacts the REND domain of both chains. Helix 3 of the THATCH domain contacts helix 5 of the REND domain. Helix 4 of the THATCH domain contacts Helix 2 of the self chain REND domain and helix 1 of the other chain.

a

b

c

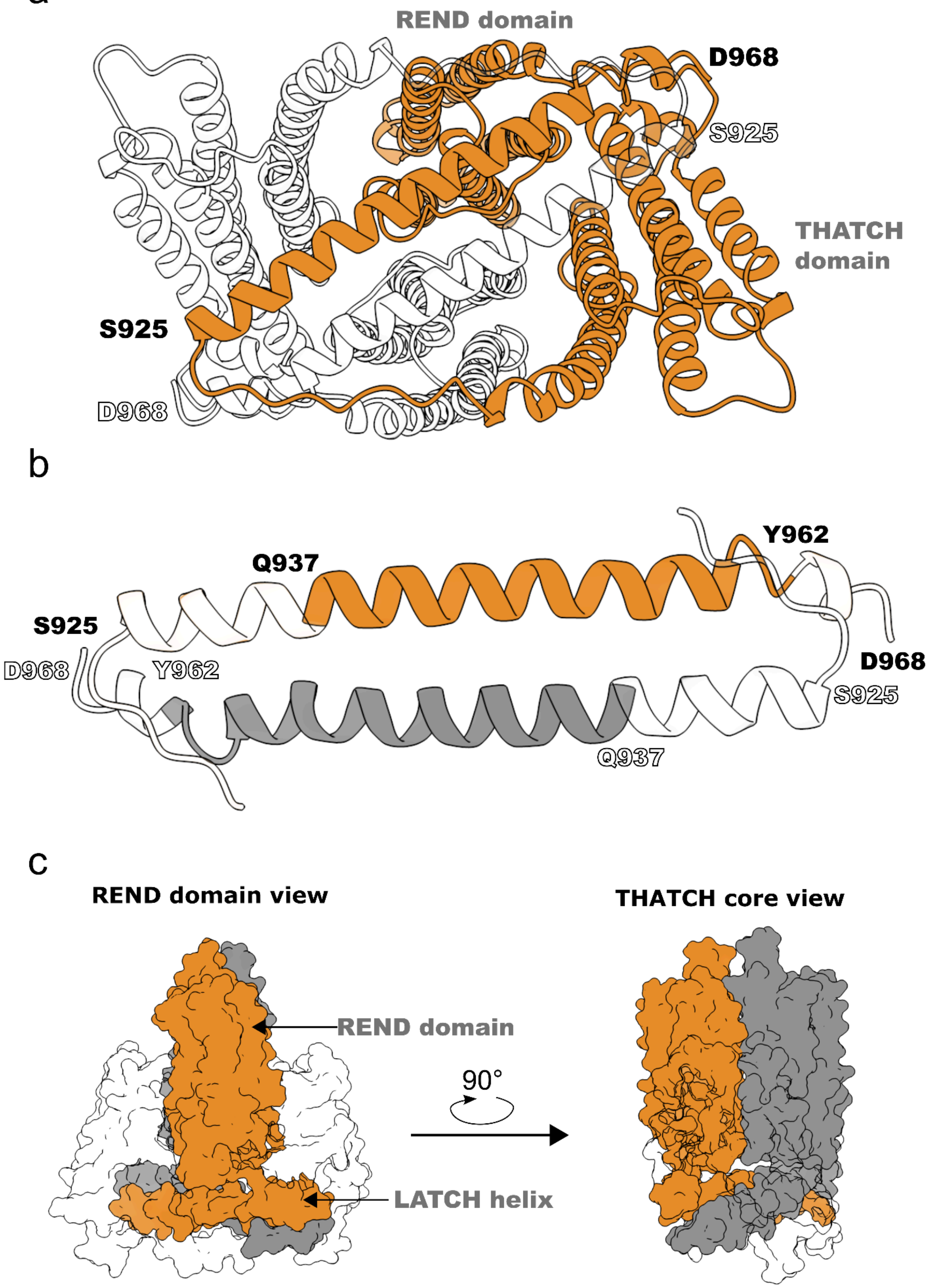


**Supplementary Figure 7: Sla2 LATCH helix forms an antiparallel dimer between the THATCH domains**

(**A**) The Sla2 cartoon highlights that the LATCH helix is antiparallel and goes from contacting the opposite chain THATCH domain to the THATCH domain of the self chain. (**B**) LATCH helix cartoon representation from our model from residues 920-968, with N- and C- terminal residues added for the LATCH Helix (925 and 968). Q937 and Y962 are labelled from the work on Talin that proposed a dimerisation motif for the THATCH domain. The antiparallel dimer proposed from homology in Talin-1 is not similar to our structure. The region 937-962 is not symmetrically aligned. The C-terminal portion of the LATCH helix contacts with not only the N-terminus of the partner helix but also the loop connecting to the THATCH core. (**C**) The surface representation of the REND and LATCH helices show that there is little buried surface area between the two domains. The REND and LATCH regions are coloured orange and grey by chain, and the THATCH core for both chains are transparent.

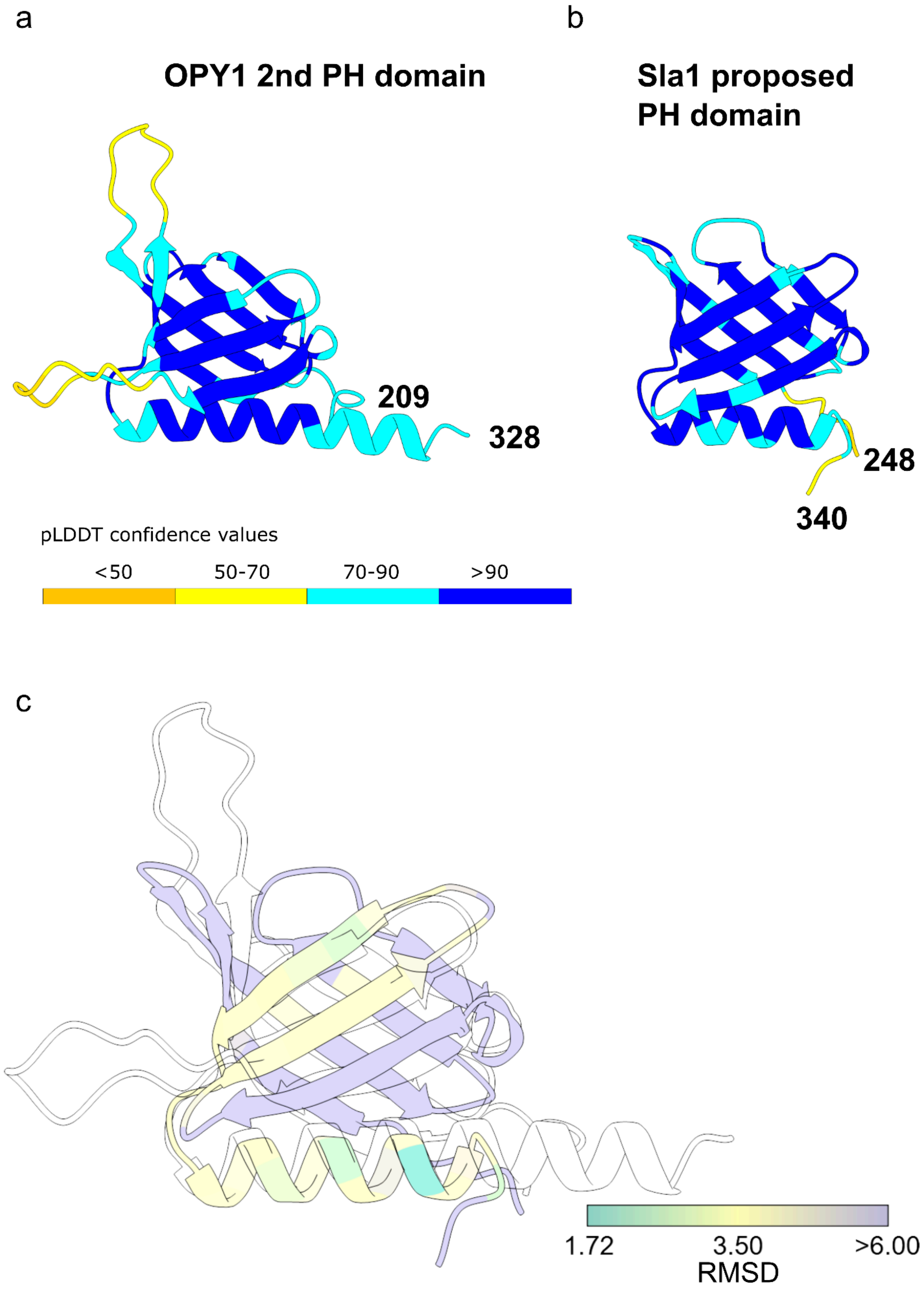


**Supplementary Figure 8: AF3 and FoldSeek elucidate a proposed PH domain in Sla1**

(**A**) AF3 model of the 2nd PH domain of OPY1 deduced from FoldSeek to be highly similar to the unknown folded region in between the SH3 domains of Sla1. (**B**) AF3 model of the folded region between the 2nd and 3rd SH3 domain of Sla1, which we now propose as an PH domain. (**C**) The proposed Sla1 PH domain aligned to OPY1 2nd PH domain as the reference structure. The Sla1 PH domain is coloured by RMSD to the reference structure. Average RMSD is 5.1 Å across the model.

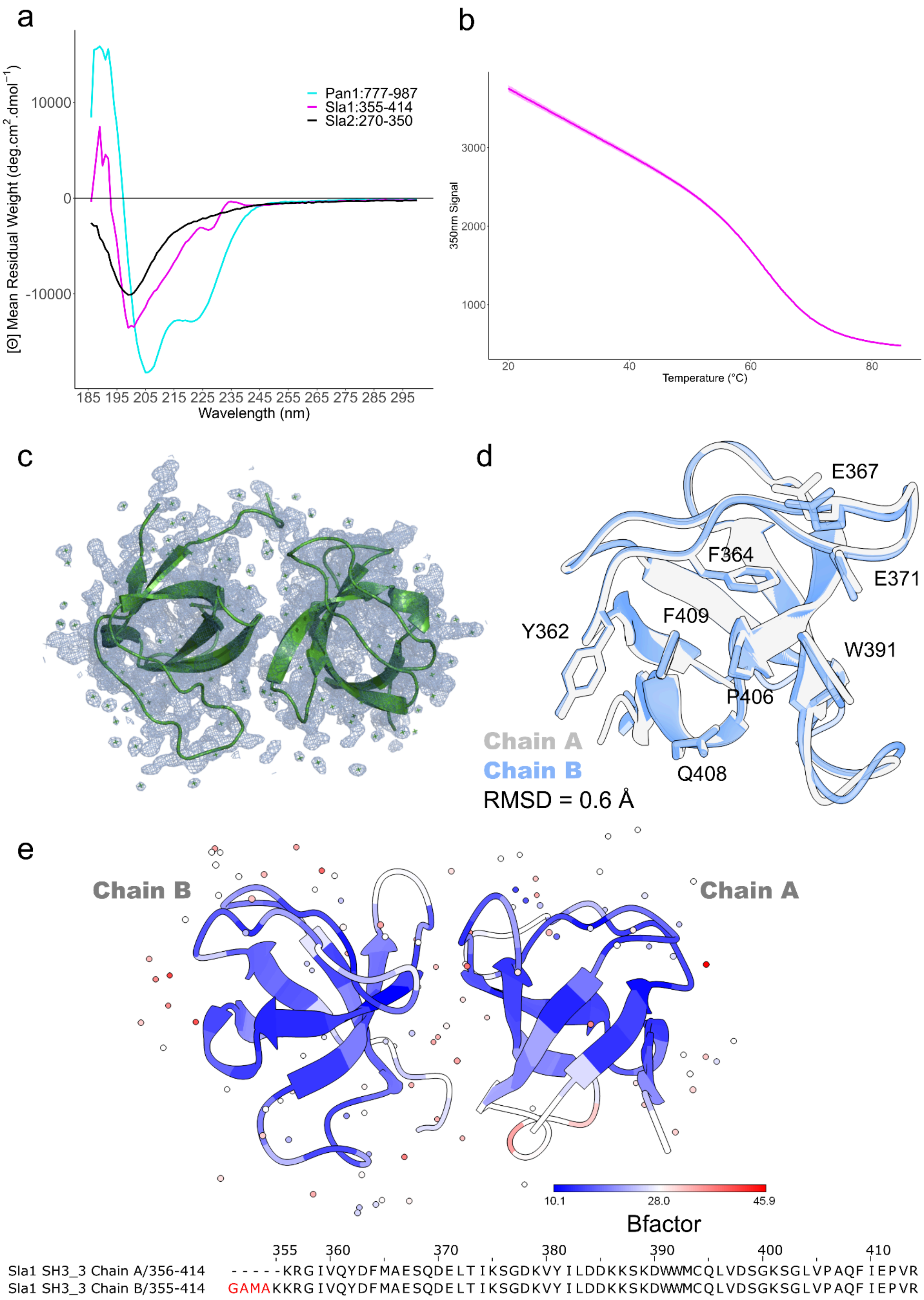

a
[Θ] Mean Residual Weight (deg.cm².dmol⁻¹)
10000
0
-10000
185 195 205 215 225 235 245 255 265 275 285 295
Wavelength (nm)
Pan1:777-987
Sla1:355-414
Sla2:270-350
b
350nm Signal
3000
2000
1000
20
40
60
80
Temperature (°C)
c
d
E367
F364
E371
Y362
F409
W391
P406
Q408
Chain A
Chain B
RMSD = 0.6 Å
e
Chain B
Chain A
10.1
28.0
45.9
Bfactor
355 360 370 380 390 400 410
Sla1 SH3_3 Chain A/356-414 - - - - -KRGIVQYDFMAESQDELTIKSGDKVYILLDDKKSKDWWMCQLVDSGKSGLVPAQFIEPVR
Sla1 SH3_3 Chain B/355-414 GAMAKKRGIVQYDFMAESQDELTIKSGDKVYILLDDKKSKDWWMCQLVDSGKSGLVPAQFIEPVR

**Supplementary Figure 9: Characterisation of Pan1:777-987, Sla2:270-350, and Sla1:355-414**

(**A**) Far-UV Circular Dichroism spectra for Pan1:777-987, Sla1:355-414, and Sla2:270-350. Secondary structure content estimations using the ChiraKit software from the eSPC online tool kit (see methods) retrieved the following percentage values for Pan1:777-987 (Alpha 33.6, Beta 10.9, Turns 14.1, Disordered 40.6); Sla1:355-414 (Alpha 13.9, Beta 28.1, Turns 11.8, Disordered 45.4) and Sla2:270-350 (Alpha 11.1, Beta 28.4, Turns 14.0, Disordered 45.0). (**B**) Thermal stability fluorescence-based assay to assess a melting curve of Sla1:355-414 at 350 nm (nanoDifferential Scanning Fluorimetry), the Tm fitted from the curve corresponds to 64.6 ± 0.1 ºC SD = 0.0067 across 3 replicates. (**C**) Models of the Sla1 SH3_3 domains, with Oxygen atoms representing the modelled water molecules, fit into the electron density map at 1 σ. (**D**) The two chains of Sla1 SH3_3 overlaid and key residues labelled from the peptide binding groove. The chains have an overall RMSD of 0.6 Å. (**E**) The complete asymmetric unit model including waters coloured by B-factor. The respective Chains are labelled as well as the sequence of residues modelled for each chain at the bottom. The average B-factor across the model is 19 Å$^2$. The sequence in black is the Sla1 SH3_3 domain sequence and the red residues are the TEV cleavage scar sequence during protein production.

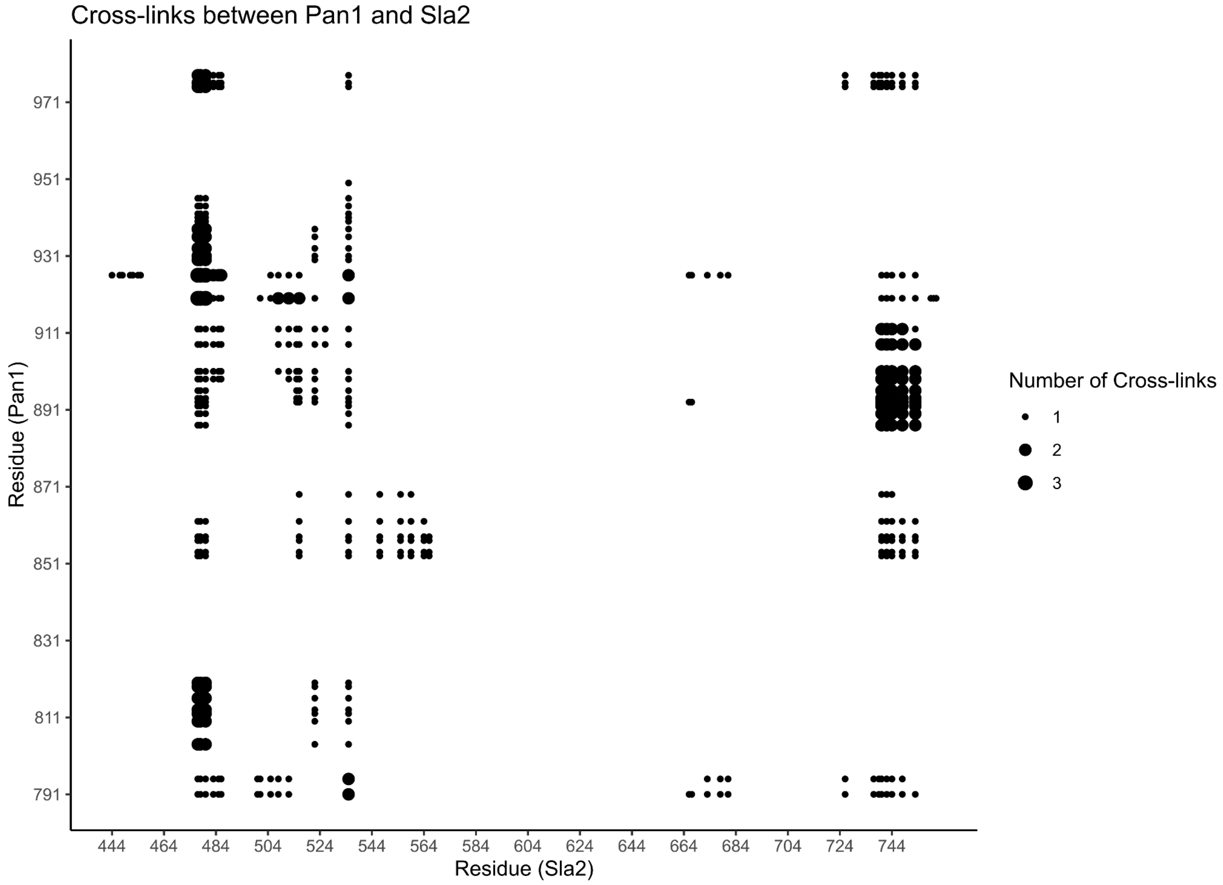


**Supplementary Figure 10: Cross-Linking MassSpectometry of Sla2 and Pan1 shows binding of Pan1 to the coiled-coil in proximity to Site 2 and Site 1**

BS3 cross-linking of Sla2:351-968 and Pan1:777-987, the cross-links were filtered to match the constructs used for MST. There were no cross-links in the N-terminal region of the Sla2 construct and there is a clustering of cross-links in the Site2/1 region. The number of cross-links in Site 2 is 3 times greater than those in the Site 1 region. There are significant cross-links in the C-terminus of the construct but this can be attributed to the flexible nature of the coiled-coil and the high number of lysines exposed on the loops of the THATCH domain.

**Supplementary Table 1: Cryo-EM data collection, refinement, and validation statistics. This table provides the parameters and statistics for the data collection, processing, refinement, and structure validation. Refinement statistics were generated using the Servalcat package in wwPDB**

| | Sla2 C-terminal region |
|---|---|
| PDB code | 9HDD |
| EMDB code | EMD-52061 |
| Data collection and processing | |
| Microscope/detector | Titan Krios G3i/K3 |
| Magnification | 120,000 x |
| Voltage | 300kV |
| Electron exposure ($e^-/Å^2$) | 45 |
| Defocus range (μm) | -0.5 to -2 |
| Pixel size (Å) | 0.68 |
| Symmetry imposed | C2 |
| Final particle images | 249299 |
| Reconstruction method | Single Particle |
| Map resolution (Å) ($FSC_{0.143}$) | 3.62 |
| Map Sharpening B factor (Å) | Local Filter |
| Refinement | |
| Initial model | AF3 model of dimeric Sla2:560-968 |
| Model composition | |

| | |
|---|---|
| **Non-hydrogen atoms** | **6296** |
| **Protein residues** | **818** |
| **R.m.s. Deviations (RMSZ)** | |
| **Bond lengths** | **0.48** |
| **Bond angles** | **0.87** |
| **Validation** | |
| **MolProbity score** | **1.2** |
| **Clashscore** | **1** |
| **Poor rotamers (%)** | **0.1** |
| **Ramachandran plot** | |
| **Favoured (%)** | **96** |
| **Allowed (%)** | **4** |
| **Disallowed (%)** | **0** |

**Supplementary Table 2: Primers used during cloning and mutagenesis of constructs:**

| Construct | Primer name | Cloning strategy | Sequence |
|---|---|---|---|
| pRS315_Sla2 | Sla2_F | SLICE | ATGTCCAGAATAGATTCAGATCTGCAGAA |
| | Sla2_R | SLICE | CTGTTATCCCTAGCGGATCCTCAATCATCATCCTGGTTATAGTAGGCATG |
| | mSc-I_F | SLICE | ACCAGGATGATGATGGCGGAGGGGGTAGCATGGTTTCTAAAGGCGAAGCCG |
| | mSc-I_R | SLICE | TATCCCTAGCGGATCCTTACTTGTACAATTCATCCATACCACCAG |
| | UTR_F | SLICE | TACCGGGCCCCCCCCCCAGCACGAAACGAAAAC |

| | | | |
|---|---|---|---|
| | UTR_F | SLICE | ATCTATTCTGGACATCCTGTTCTAGCTGCTAGTACTATCACTACTACTGCTATG |
| Sla2_S1 | Sla2_S1 | Knock_out | CAGCCATAGCAGTAGTAGTGATAGTACTAGCAGCTAGAACAGGATGCGTACGCTGCAGGTCGAC |
| Sla2_S2 | Sla2_S2 | Knock_out | ATATATTTATATTAACGTTTATCTTTATATATAAAAAGTACAATTCATGATCAATCGATGAATTCGAGCTCG |
| pETM30-Sla2cc/pRS315 Sla2_deltaYYR | deltaYYR_F | QuikChange | GGATCAATTGGATGTTTGGGAAAGAAAAGCTGAGTCTTTAGCCAAGCTA |
| | deltaYYR_R | QuikChange | TAGCTTGGCTAAAGACTCAGCTTTTCTTTCCCAAACATCCAATTGATCC |
| | deltaYYR_F2 | QuikChange | GAAAAGCTGAGTCTTTAGCCAAGCTAGCCTCCCAGTTGGCTCAAGAGCATCTAAATCTTTTAC |
| | deltaYYR_R2 | QuikChange | GTAAAAGATTTAGATGCTCTTGAGCCAACTGGGAGGCTAGCTTGGCTAAAGACTCAGCTTTTC |
| pETM30-Sla2cc/pRS315 Sla2_deltaMotif | deltaMotif_F | QuikChange | GCTGCACCTGGTTTTGATGAAGCGCAGTTAAAGGTGAATAGTGCGCAGGA |
| | deltaMotif_R | QuikChange | CCAGGTGCAGCACGTGCAGCCTCTTGACGCAACTGGGAGTATAGC |
| pETM30-Sla2cc/pRS315 Sla2_delta515-546 | delta515-546_F | QuikChange | GAATAGTGCGCAGGAATCCATTCAGTCCATTAATAATGCAGAGGCGGAC |
| | delta515-546_R | QuikChange | GTCCGCCTCTGCATTATTAATGGACTGAATGGATTCCTGCGCACTATTC |
| pETM30_Sla2_IDR | IDR_F | SLICE | TTTTCAGGGCGCCATGGCCGTGGACGAGTCAAAAGAGATTAAG |
| | IDR_R | SLICE | GCTCGAATTCGGATCCTTACTGTGGAAAAATGGCGTTAG |
| pETM30_CLC_1-80 | CLC_1-80_F | QuikChange | GTGCGGTGAGCAGCGATTGACTTAAGCAATTGGGATCCTAAT |
| | CLC_1-80_R | QuikChange | ATTAGGATCCCAATTGCTTAAGTCAATCGCTGCTCACCGCAC |
| pETM30_CLC_70-140/pETM30_CLC_70-233 | CLC_70_F | SLICE | TTTCAGGGCGCCATGATTAACAGCGCGAACGGTG |

| | | | |
|---|---|---|---|
| pETM30_CLC_70-140 | CLC_140_R | SLICE | GAATTCGGATCCTTACTCGTCCTTCAGATCTTTTTCGTG |
| pETM30_CLC_70-233 | CLC_233-R | SLICE | GAATTCGGATCCTTACGCGCCCGGCGC |
| pnEA-vHisGST_CLC_351-968 | Sla2_351_F | SLICE | TTTACTTCCAGGGCCATATGGCGACGGCACAAATGCAG |
| | Sla2_R | SLICE | AGACTATTAGGATCCATCATCATCCTGGTTATAGTAGGCATGC |
| pETM30_Sla1_SH3_3 | Sla1_F | SLICE | TTCAGGGCGCCATGGCCAAGAAAAGGGGAATAGTACAATATG |
| | Sla1_R | SLICE | CTCGAATTCGGATCCTTAACGAACCGGCTCGATGAAC |
| pETM30_Pan1_777-987 | Pan1_F | SLICE | TTCAGGGCGCCATGGCCGCGAAACCAAAATATGCTGGG |
| | Pan1_R | SLICE | CTCGAATTCGGATCCTTAACAAATAAATCAGTGACTGAGTCATCTCCC |

**Supplementary Table 3: X-ray crystallography data collection, refinement, and validation statistics. This table provides the parameters and statistics for the data collection, processing, refinement, and structure validation. Refinement statistics were generated using the Servalcat package in wwPDB. The values in parentheses refer to the highest resolution shell.**

| | **Sla1 SH3_3** |
|---|---|
| **PDB ID** | **9HDB** |
| **Data collection** | |
| **Beamline** | **PETRA III / P13** |
| **Space Group** | **P 21 21 21** |
| **Cell Dimensions** | |
| **a , b, c (Å)** | **38.53, 50.42, 51.97** |
| **α, β, γ (°)** | **90.00, 90.00, 90.00** |
| **Resolution (Å)** | **30.95-1.491 (1.544 - 1.491)** |

| | |
|---|---|
| Rpim | 0.02661 (0.136) |
| <I/σI> | 3.29 (at 1.49Å) |
| Mean (I/sd(I)) | 15.6 (4.8) |
| CC 1/2 | 0.998 (0.962) |
| Completeness (%) | 99.9 % (99.9 %) |
| Redundancy | 10.0 (10.0) |
| Refinement | |
| No. of reflections (work/free) | 17084/847 |
| Rwork/Rfree | 0.142/0.189 |
| Ramachandran favoured regions (%) | 99.16 |
| Ramachandran allowed regions (%) | 0 |
| Ramachandran outliers (%) | 0.84 |
| Rotamer outliers (%) | 0.0 |
| Clashscore | 0.51 |
| No. of non-Hydrogen atoms | |
| Protein | 1099 |
| Water | 111 |
| B-factors ($Å^2$) (Average) | 19.48 |
| Protein | 18.27 |
| Water | 30.28 |
| RMS deviations | |
| Bond lengths (Å) | 0.017 |
| Bond angles (°) | 2.02 |
